# High throughput screening identifies broad-spectrum Coronavirus entry inhibitors

**DOI:** 10.1101/2023.12.04.569985

**Authors:** Suman Khan, Efrat Ozer Partuk, Jeanne Chiaravalli, Noga Kozer, Khriesto A. Shurrush, Yael Elbaz-Alon, Nadav Scher, Emilie Giraud, Jaouen Tran-Rajau, Fabrice Agou, Haim Michael Barr, Ori Avinoam

## Abstract

The Covid-19 pandemic highlighted the pressing need for antiviral therapeutics capable of mitigating infection and spread of emerging coronaviruses (CoVs). A promising therapeutic strategy lies in inhibiting viral entry mediated by the Spike (S) glycoprotein. To identify small molecule inhibitors that block entry downstream of receptor binding, we established a high-throughput screening (HTS) platform based on pseudoviruses. We employed a three-step process to screen nearly 200,000 small molecules. First, we identified potential inhibitors by assessing their ability to inhibit pseudoviruses bearing the SARS-CoV-2 S glycoprotein. Subsequent counter-screening against pseudoviruses with the Vesicular Stomatitis Virus glycoprotein (VSV-G), yielding sixty-five SARS-CoV-2 S-specific inhibitors. These were further tested against pseudoviruses bearing the MERS-CoV S glycoprotein, which uses a different receptor. Out of these, five compounds including the known broad-spectrum inhibitor Nafamostat, were subjected to further validation and tested them against pseudoviruses bearing the S glycoprotein of the alpha, delta, and omicron variants as well as against *bona fide* SARS-CoV-2 *in vitro*. This rigorous approach revealed a novel inhibitor and its derivative as a potential broad-spectrum antiviral. These results validate the HTS platform and set the stage for lead optimization and future pre-clinical, *in vivo* studies.

## Introduction

Coronaviruses (CoVs) have garnered global attention due to their potential for causing severe diseases in humans. The most notable among these are SARS-CoV, MERS-CoV, and SARS-CoV-2, each responsible for significant disease outbreaks^1^. As zoonotic pathogens, CoVs continue to pose a constant threat to global health due to the potential for cross-species transmission, underscoring the need for broad-spectrum antiviral inhibitors.

The viral Spike (S) glycoprotein of CoVs mediates fusion of the viral envelope with the host cell membrane, which is essential for infection and delivery of the viral genetic material into host cells^2–4^. This process is conserved across all coronaviruses, positioning the S glycoprotein as a promising target for broad-spectrum antiviral strategies^5,6^. The S glycoprotein is a class I viral fusogens, comprised of two subunits: S1, involved in host cell recognition and binding, and S2, which mediates membrane fusion^3,7^.

Current therapies for CoV infection largely aim to disrupt the S1 domain-mediated host recognition^8^. However, these strategies face significant limitations, particularly with the emergence of SARS-CoV-2 variants carrying mutations in the S1 domain that enhance receptor binding and facilitate immune evasion ^9–12^. Additionally, the variability in the cellular receptors recognized by different CoVs presents a challenge for achieving broad inhibition with S1 domain-targeted strategies. For instance, SARS-CoV and SARS-CoV-2 recognize angiotensin-converting enzyme 2 (ACE2)^13,14^, while MERS-CoV interacts with dipeptidyl peptidase 4 (DPP4)^15^. Further, the identification of new SARS-CoV-2 receptors, such as TMEM106B^16^ highlights the adaptability of CoVs in exploring alternative receptors.

Strategies to inhibit host proteases like Transmembrane protease serine 2 (TMPRSS2)^17–19^ and Cathepsins^20,21^, which facilitate CoV entry by cleaving the S protein and activating the fusogenic activity of the S2 domain, have also been explored. Protease inhibitors like Nafamostat^22^ and Camostat^19^ have demonstrated some efficacy against multiple CoVs^23,24^. However, their inhibition spectrum remains uncertain due to the adaptability of CoVs in exploring alternative proteases^25^, and their evolution to include a polybasic furin cleavage site^26,27^, thereby limiting the strategy of targeting a single protease for broad-spectrum inhibition.

In contrast, the S2 domain presents as a promising target for managing CoV infection. Cross-reactive neutralizing antibodies (nAbs) against the S2 domain have been identified in individuals who have not contracted SARS-CoV-2, as well as patients infected with various CoVs^28,29^. This compelling evidence is reinforced by the essential role of the S2 domain in the universally conserved biophysical process of membrane fusion. Additionally, in comparison to the S1 domain, the S2 domain has exhibited lower mutation rates in emerging SARS-CoV-2 variants, which is further supported by phylogenetic analyses showing a higher degree of sequence conservation in the S2 domains of diverse CoV clades^12,30–32^. These characteristics of the S2 domain suggest its potential as a broad-spectrum therapeutic target. However, targeting the S2 domain is challenging because it cannot be expressed independently of the S1 domain. Hence, current FDA-approved drugs for treating COVID-19 patients, such as Remdesivir^33^, Molnupiravir^34^ and Nirmatrelvir^35^, do not specifically target the S2 domain.

High throughput screening (HTS) has been used to identify antiviral leads for various viruses^36–38^. With the emergence of SARS-CoV-2, several HTS assays were swiftly developed, predominately focusing on FDA-approved drugs, to reduce the development time by re-purposing existing drugs^39–41^. Despite numerous efforts, few novel and efficient antivirals were identified^42–45^. Furthermore, while numerous *in silico* and *in vitro* HTS approaches targeting viral entry or viral replication have been developed, efforts specifically dedicated to the identification of broad-spectrum CoV antivirals through HTS have been sparse^46,47^.

To address this gap, we adopted a pseudotyped Vesicular Stomatitis Virus (VSV) model^48^ permitted robust quantification of CoV Spike glycoprotein mediated infection. We established an HTS platform and screened approximately 200,000 diverse chemical compounds. We targeted the S2 domain of the S glycoproteins by screening against the glycoproteins of two distinct CoVs that bind different cellular receptors^30,49–51^. Our extensive HTS efforts resulted in the identification and validation of a novel broad-spectrum antiviral compound.

## Results

### Pseudoviruses expressing fluorescent reporters enable robust infection quantification

To develop an effective high-throughput screening (HTS) assay for SARS-CoV-2, we produced pseudoviruses featuring the S glycoprotein of the SARS-CoV-2 Wuhan variant (VSVΔG-S_W_) on a VSV backbone lacking VSV-G (VSVΔG) **(Fig. 1 A)**. We chose to work with VSVΔG which express a fluorescent reporter from the viral genome after infection **(Fig. 1 B)**. We employed a fluorescent reporter, instead of the more commonly used luciferase, because it allows direct visualization and robust quantification of single infection events, overcoming the need for averaging and the additional processing steps for the Luciferase enzymatic reaction^52^ **(Fig. 1 C)**.

**Fig. 1.**
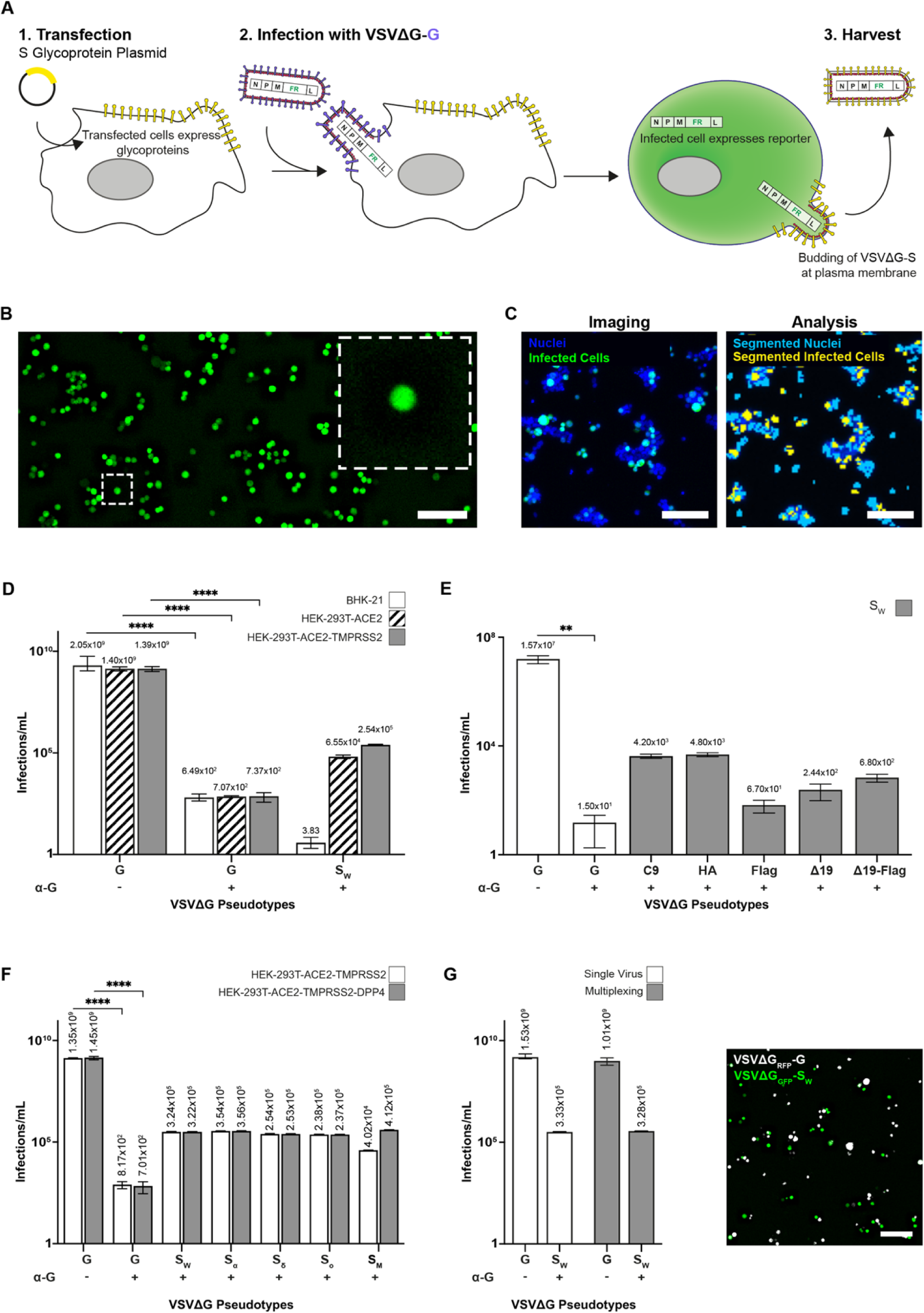
Production of high titer VSVΔG pseudovirus and quantification of single infection events. **(A)** A schematic representation of the glycoprotein complemented VSVΔG pseudoviruses producing process. Viral glycoproteins (yellow) are overexpressed on cell membranes using a plasmid encoding the chosen glycoproteins. These transfected cells are then infected with VSVΔG-G, resulting in viral-induced expression of a fluorescent reporter (FR) and VSV structural proteins that assemble on the cell surface, incorporating the desired glycoproteins into the VSVΔG pseudovirus. **(B)** A widefield image of cells infected with VSVΔG pseudoviruses expressing a fluorescent reporter (GFP; green). Infected cells become round after infection due to the virus-induced Cytopathic Effect. **(C)** A high magnification overlay image showing the infected GFP-positive cells (green) and total nuclei (blue), and the respective segmentation showing infected cells (yellow) and total nuclei (cyan). **(D-G)** Pseudoviral titers in infections/ml. An antibody against VSV-G (α-G) was utilized to neutralize residual VSVΔG-G infection from production, ensuring accurate titration of the heterologous pseudoviruses. **(D)** Titer of VSVΔG-S_W_ showing enhanced infection of cells expressing the innate receptor, ACE2 and host protease, TMPRSS2. **(E)** Titer of VSVΔG-S_W_ showing the effect of modifications to the cytosolic tail of S_W_. **(F)** Titer of VSVΔG-G, Wuhan (S_W_), Alpha (S_α_), Delta (S_δ_), Omicron (S_ο_), and MERS-CoV S (S_M_) showing similar infection levels in HEK-293T cells expressing ACE2-TMPRSS2, with and without DPP4. **(D-F)** VSVΔG-G Pseudoviruses infected all cell lines at similar levels. **(G)** Pseudoviral infections/ml of VSVΔG_RFP_-G and VSVΔG_GFP_-S_W_ separately or simultaneously results in equivalent infection rates. (Right) A high magnification overlay image of a well showing VSVΔG_RFP_-G and VSVΔG_GFP_-S_W_ infected cells (White and green, respectively). The statistical significance of antibody activity was also determined. P: **≤0.01, ****≤0.0001 (two-tailed unpaired t-tests). N(experiments)=3, n(readings)=9. Error bars represent the SEM. Scale bar is 100 μM (B, C and G).

We confirmed the specific tropism of the VSVΔG-S_W_ pseudoviruses by comparing infected ACE2-overexpressing Human Embryonic Kidney (HEK-293T-ACE2) cells and ACE-2-deficient Baby Hamster Kidney (BHK-21) cells **(Fig. 1 D)**. To eliminate potential residual infections from VSVΔG-G that might have been left over during production, we performed all experiments in the presence of a neutralizing antibody against VSV-G. We observed 10,000-fold more infections in HEK-293T-ACE2, with an additional 3.8-fold increase in cells co-expressing TMPRSS2 (HEK-293T-ACE2-TMPRSS2), consistent with the tropism of SARS-CoV-2 **(Fig. 1 D)**.

We then explored methods to enhance virus titers, comparing modifications to the cytosolic tail of the S glycoprotein and optimizing the production, and infection procedures **(Fig. 1 E and S1 A-B)**. Modifications included truncating the 19-amino acid ER retention sequence^53,54^ and adding an HA, flag or C9 tag to the C-terminus of the S glycoprotein^19,55^. Optimal titers were achieved with C9 and HA tagged S glycoproteins without further modifications **(Fig. 1 E)**. Moreover, we obtained peak titers when the cell supernatant containing VSVΔG-S_W_ was harvested 30 hours after infection with the VSVΔG-G helper virus. Notably, infection rates doubled when cultures were centrifuged post-infection **(Fig. S1 A-B)**. These optimizations significantly improved the sensitivity and reproducibility of infection counts.

Subsequently, we produced and obtained high titer pseudoviruses featuring the S glycoproteins of MERS-CoV (VSVΔG-S_M_) and SARS-CoV-2 variants Alpha (VSVΔG-S_α_), Delta (VSVΔG-S_O_), and Omicron (VSVΔG-S_ο_), and quantified infection rates in both HEK-293T-ACE2-TMPRSS2-DPP4 and HEK-293T-ACE2-TMPRSS2 cell lines **(Fig. 1 F)**. Importantly, we assessed the infection rates of VSVΔG pseudoviruses expressing either GFP or RFP and featuring the S glycoprotein (VSVΔG_GFP_-S_W_) or VSV-G (VSVΔG_RFP_-G) respectively **(Fig. 1 G)**. Our results showed that comparable titers were achieved in both single and multiplexed infections, demonstrating the feasibility of multiplexing the assay using pseudoviruses expressing different fluorophores **(Fig. 1 G)**.

### Optimization and validation of a high throughput screening assay for infection inhibitors

Building on the robust segmentation and quantification capabilities of our automated imaging system and the fluorescence-based assay using VSVΔG pseudoviruses **(Fig. 1 B-C)**, we tailored the assay for high throughput screening (HTS) in a 384-well format **(Fig. 2 A)**. To streamline the plating process and ensure robust and reproducible quantification, compounds in DMSO were pre-plated in 384-well plates. Then, 10 µl of pseudovirus suspension was added to achieve 500-1000 infections per well. Subsequently, 10,000 HEK-293T-ACE2-TMPRSS2 cells were added to each well and the plates were centrifuged and incubated for 24 hours **(Fig. 2 A)**.

**Fig. 2.**
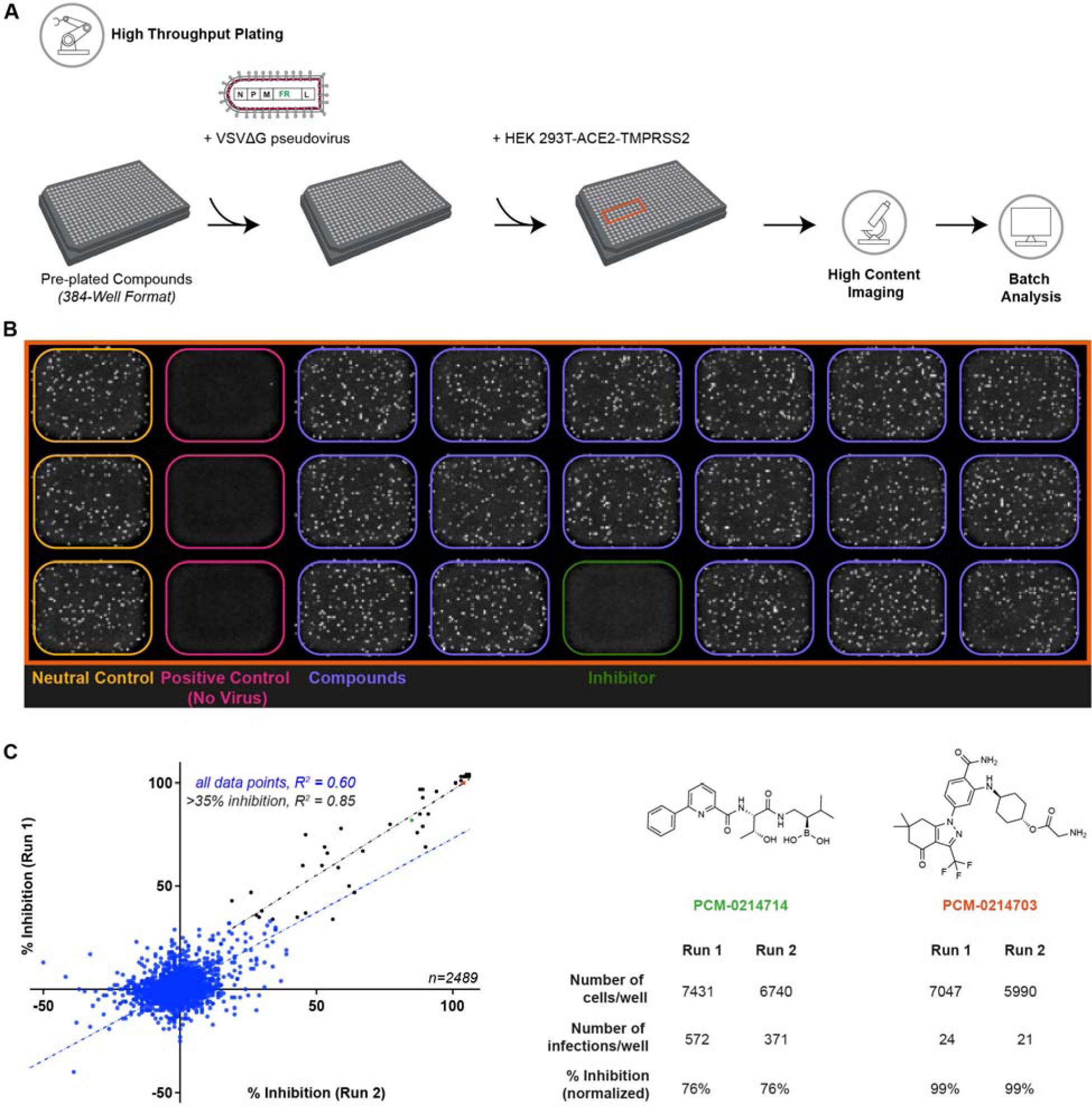
The pseudovirus-based HTS platform demonstrates high reproducibility. **(A)** Schematic of the high content screening pipeline: plating, imaging, and analysis. Compounds were pre-plated, and pseudoviruses, pre-incubated with α-G to neutralize any residual VSVΔG-G infection, were added in individual wells before introducing HEK-293T cells stably overexpressing SARS-CoV-2 receptor ACE2 and protease TMPRSS2. After 24 hours, the nuclei were stained, and the plates were imaged. Images were subsequently segmented and quantified **(B)** Overview image of a portion of a 384-well screening plate (orange box in A). The neutral control (yellow wells) indicates 100% infection, and the positive control (magenta wells) signifies 0% infection or 100% inhibition. The green well depict an example of a compound with ∼90% inhibition. **(C)** Scatter plot of 2489 compounds tested as single point, on two different days. Blue line is the fit for the compounds that inhibited >35%. The inhibitions were reproducible with a high correlation fit (R^2^=0.85). Black line represents the fit for the rest of the compounds (R^2^=0.60). (Right) Examples of two inhibitors showing similar inhibition values after normalization and despite having different total cell and infection counts.

Post infection, the nuclei were stained to estimate cell count. Automated HTS imaging was then used to obtain images of the infected cells and nuclei in each well. These images were automatically segmented and analyzed by a dedicated pipeline **(Fig. 1 C and 2 B)**. The first columns of each plate, which contained viruses and cells, was used as a neutral control, providing the baselines for the infection and cell number counts **(Fig 2 B and S2 A)**. The raw infection and cell number counts from each well were normalized with the geometric means of the infection and cell number counts in the neutral control to determine the baselines for the inhibition and cytotoxicity profiles for each compound **(Fig. 2 B and S2 A)**. The second columns contained only cells and served as the baseline for 0% infection as the geometric mean of counts in these wells, which also served as the positive controls **(Fig. 2 B and S2 A)**. For consistency, all wells without compounds were supplemented with DMSO to achieve a final concentration of 0.1%.

To test the reproducibility of the assay and ascertain if a one-time screen at a single concentration for each compound would be likely to identify putative inhibitors, we ran a pilot screen with a subset of 2,489 compounds, approximately 1.25% of the entire compound library **(Fig. 2 C)**. Each compound was assessed at a concentration of 10 μM for its ability to inhibit VSVΔG-S_W_ in two separate experiments. Compounds demonstrating at least 35% inhibition were reliably detected with similar inhibition levels, independent of day-to-day variations and regardless of the number of cells or the raw infection counts, confirming the robustness of the assay **(Fig. 2 C)**.

### A three-tiered screen identifies potential broad spectrum S2-domain inhibitors

To identify potential S2-domain specific inhibitors, we divided our screening process into three distinct tiers **(Fig 3 A)**. The primary screen evaluated an extensive library of approximately 200,000 compounds against VSVΔG-S_W_ at a single concentration of 10 μM. This process identified 733 compounds capable of inhibiting VSVΔG-S_W_ infection by at least 35% while maintaining cell viability of 65% (Supplementary Table 1). For the secondary screen, the 733 compounds were tested against VSVΔG-G to distinguish S_W_-specific inhibitors from non-specific ones. Since VSVΔG-S_W_ and VSVΔG-G share the VSVΔG backbone and only differ in the surface glycoproteins, compounds inhibiting VSVΔG-S_W_ but not VSVΔG-G were deemed S_W_-specific. The screening identified 65 Spike-specific inhibitors that inhibited VSVΔG-S_W_ infection by at least 35% without inhibiting VSVΔG-G infection more than 35% (Supplementary Table 2).

**Fig. 3.**
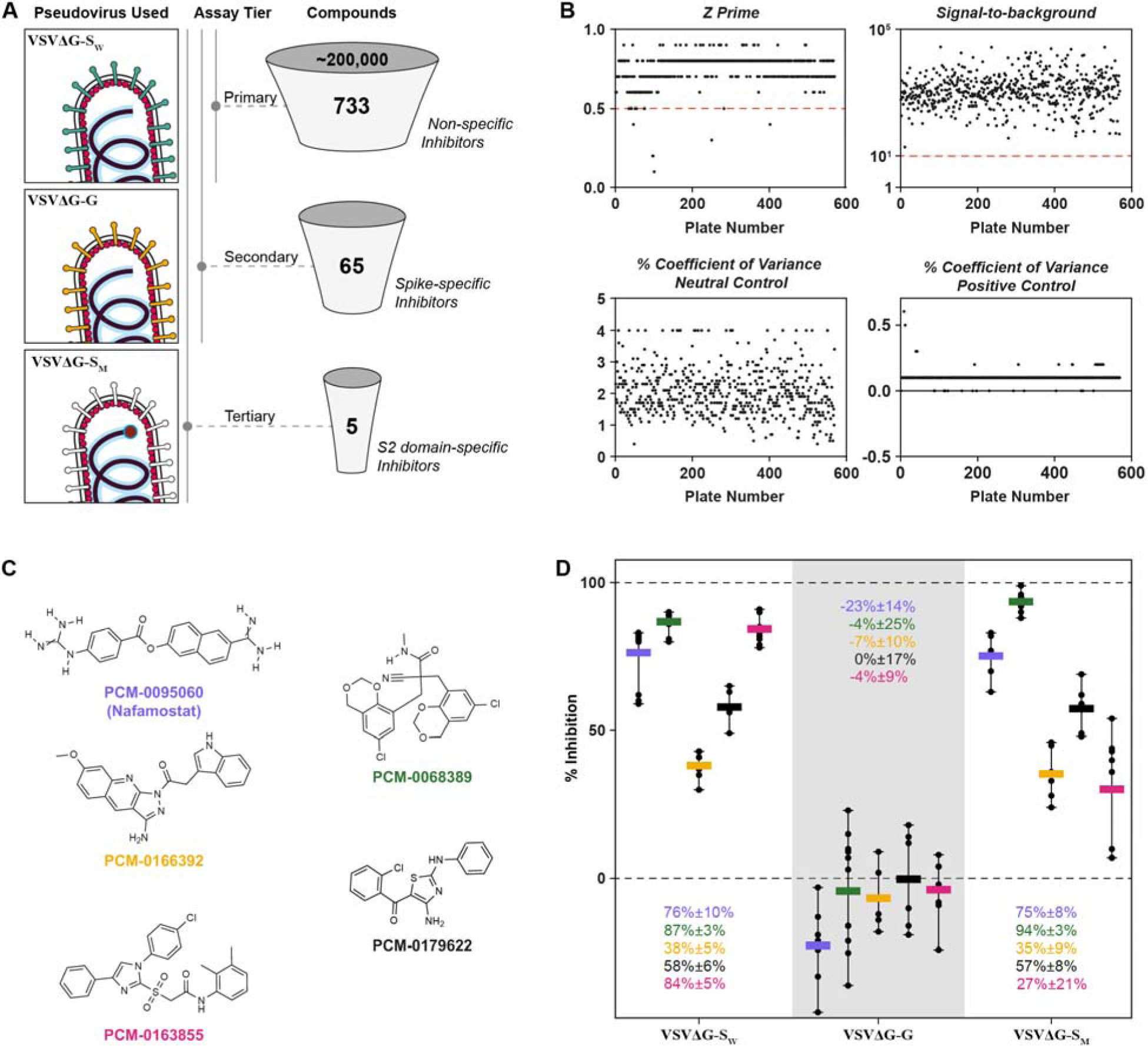
A three-tiered screen identifies putative entry inhibitors of CoVs. **(A)** Schematic showing primary screening of ∼200,000 compounds against Wuhan SARS-CoV-2 Spike (VSVΔG-S_W_) yielded 733 putative Spike-specific and non-specific inhibitors. S_W_ specific inhibitors were identified by a secondary screen against VSV-G (VSVΔG-G) yielding 65 putative inhibitors. The tertiary screen with MERS-CoV Spike (VSVΔG-S_M_) and initial validation resulted in 5 compounds that were putatively broad-spectrum inhibitors. **(B)** Plots showing HTS parameters to determine the robustness of our screen: Z’ factor, signal-to-background, percentage of coefficient of variance for positive and negative for the primary screen. Red line denotes the cut off for a robust plate. Five plates that failed to meet the cut-off for Z prime were manually checked for data quality. **(C)** The chemical structures of the four novel compounds and Nafamostat that were commercially resourced. **(D)** Plot of the inhibition of the resourced compounds against the three viruses. All the compounds selectively inhibit SARS-CoV-2 and MERS-CoV Spikes without inhibiting VSV-G. Error bars represent the range. Values represent mean inhibitions ± standard deviations.

Subsequently, these 65 compounds were subjected to a tertiary screen against VSVΔG-S_M_, to segregate S2-specific inhibitors from receptor binding inhibitors. Since MERS-CoV utilizes DPP4 as its host receptor, compounds inhibiting both VSVΔG-S_W_ and VSVΔG-S_M_ are likely S2-specific. This screen yielded 22 putative S2-domain specific inhibitors that reduced VSVΔG-S_M_ infection by at least 35% (Supplementary Table 3). Out of these, we chose the 11 most drug-like inhibitors that were previously unreported, and Nafamostat, a known TMPRSS2 protease inhibitor, which we used as a positive control in subsequent experiments^22,56^.

To evaluate the screening platform, we calculated HTS parameters such as Z’ factor, signal-to-background (S/B), and coefficient of variation (%CV) for 570 plates from all three screening levels^69,70^ (**Fig. 3 B)**. These showed the robust performance of the screening platform with a high Z’ factor (0.83±0.5) and a very high S/B (10^3^), suggesting excellent sensitivity and accuracy. In addition, the low %CV (2.5±6% for neutral controls and 0.1±3% for positive control) indicates the high reproducibility and precision of the platform. Plates that failed to meet the accepted Z’ cut-off of 0.5 were manually checked and validated before data normalization.

To ensure the reliability of the HTS platform and rule out potential false positives due to compound degradation or quality, we resourced 4 of the 11 novel compounds (PCM-0068389, PCM-0166392, PCM-0179622, PCM-016855; Supplementary Table 3) and Nafamostat that were available from alternative vendors and reassessed their activity against VSVΔG-S_W_ VSVΔG-G, and VSVΔG-S_M_, **(Fig. 3 C-D)**. These compounds demonstrated varied inhibitory activity against VSVΔG-S_W_ (38%-87%) and VSVΔG-S_M_ (27%-94%) but did not significantly inhibit VSVΔG-G (−23% to 0%) confirming their selectivity **(Fig. 3 D)**.

### Dose response and cytotoxicity of putative candidate compounds

Next, we evaluated the cytotoxicity (CC_50_) and IC_50_ value for each compound using a concentration range of 0.3125 - 40 μM against VSVΔG-S_W_, VSVΔG-G, and VSVΔG-S_M_, as well as VSVΔG-S_α_, VSVΔG-S_O_, and VSVΔG-S_ο_. Since the purity of these compounds ranged from 70% to 95%, and as an additional validation step, we performed these experiments on the compounds before and after HPLC purification. All compounds showed low to moderate cytotoxicity before and after purification **(Fig. S3 and 4** respectively).

**Fig. 4.**
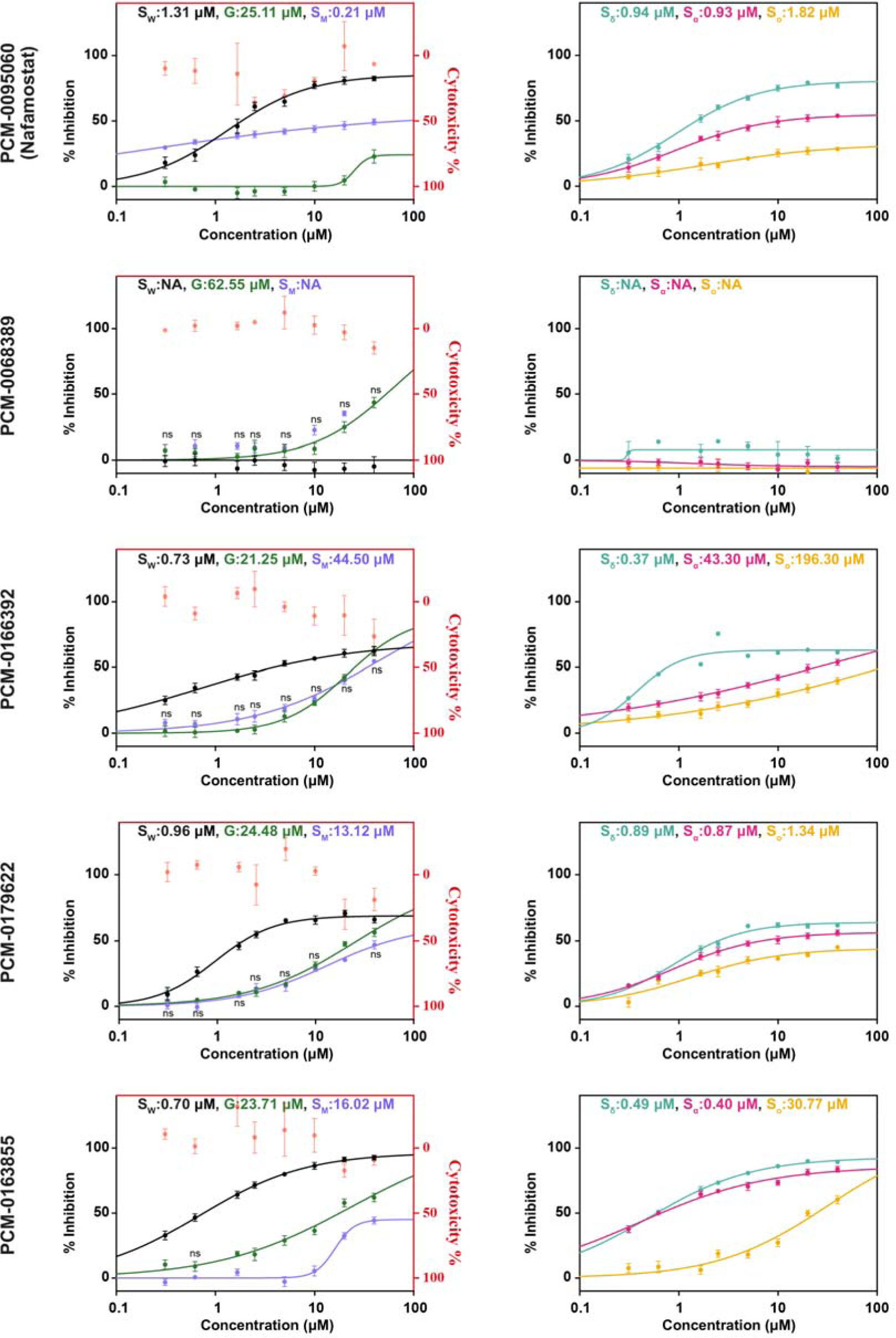
Dose-response activity and cytotoxicity of HPLC-purified compounds. **(Left)** Dose-response plot of the purified candidates against VSVΔG-S_W_, VSVΔG-S_M_ or VSVΔG-G and their cytotoxity profile. **(Right)** Dose-response plots of hits against pseudoviruses with glycoproteins of SARS-CoV-2 variants (VSVΔG-S_α_: Alpha, VSVΔG-S_O_: Delta, VSVΔG-S_ο_: Omicron). Dose-response curve were fitted with a variable slope (four-parameter logistic model). Error bars represent the SEM. ns are the readings where there is no statistically significant difference between VSVΔG-S_M_ and VSVΔG-G at a given concentration. For all other readings, the P<0.05 (two-tailed unpaired t-tests).

Nafamostat showed similar IC_50_ values against VSVΔG-S_W_ (0.90 μM vs 1.31 μM) and VSVΔG-S_M_ (0.13 μM vs 0.21 μM) before and after purification, with no significant activity against VSVΔG-G, and was also active against VSVΔG-S (0.92 μM vs 0.93 μM), VSVΔG-S_O_ (0.31 μM vs 0.94 μM), and VSVΔG-S_ο_, but the latter only after purification (Not calculated vs 1.81 μM) (**Fig. S3 and 4)**. PCM-0068389, while showing extremely promising activity and selectivity before purification, lost all activity against VSVΔG-S_W_, its variants and VSVΔG-S_M_ after purification and showed some activity against VSVΔG-G at concentrations higher than 10 μM **(Fig. S3 and 4)**. PCM-0166392 and PCM-0179622 while retaining relatively high activity against VSVΔG-S_W_ and its variants were not selective against VSVΔG-S_M_ yielding dose-response curves that were not significantly different from VSVΔG-G (**Fig. S3 and 4)**.

PCM-0163855 retained high selectivity to VSVΔG-S_W_ (0.60 μM vs 0.70 μM), VSVΔG-S_α_ (0.37 μM vs 0.40 μM), and VSVΔG-S_O_ (0.45 μM vs 0.49 μM), with only moderate activity against VSVΔG-S_ο_ (41.18 μM vs 30.77 μM), but was inactive against VSVΔG-S_M_, showing no significant difference from VSVΔG-G above 10 μM concentration **(Fig. S3 and 4)**. Consistently, PCM-016855 inhibited the replication of *bona fide* SARS-CoV-2 Delta (B.1.617.2) in VeroE6 cells with and without TMPRSS2 overexpression (6.71 μM and 6.43 μM respectively; **Fig. S4 A**).

### A PCM-016855 derivative, is a broad-spectrum CoV inhibitor

Following up on these promising results, we synthesized PCM-0163855 and its sulfoxide derivative PCM-0282478 (**Fig. S5**). We evaluated their IC_50_ value against VSVΔG-S_O_, VSVΔG-S_ο_, VSVΔG-G, and VSVΔG-S_M_, and *bona fide* SARS-CoV-2 variants, Delta (B.1.617.2), XBB.1.5 and CH.1.1 **(Fig. 5 A and Fig. S6 A)**. Dose-response curves demonstrated that PCM-0282478 was approximately 10 times more potent than synthesized PCM-0163855 against VSVΔG-S_O_ (1.13 μM and 14.40 μM respectively) and selective against VSVΔG-S_ο_ and VSVΔG-S_M_ (14.67 μM and 17.12 μM respectively). Neither PCM-0163855 nor PCM-0282478 inhibited VSVΔG-G up to 10 μM. Furthermore, only PCM-0282478 inhibited *bona fide* SARS-CoV-2 Delta (B.1.617.2) replication in VeroE6 cells (4.49 μM; **Fig. 5 A**), and neither inhibited XBB.1.5 and CH.1.1 at the concentration range tested.

**Fig. 5.**
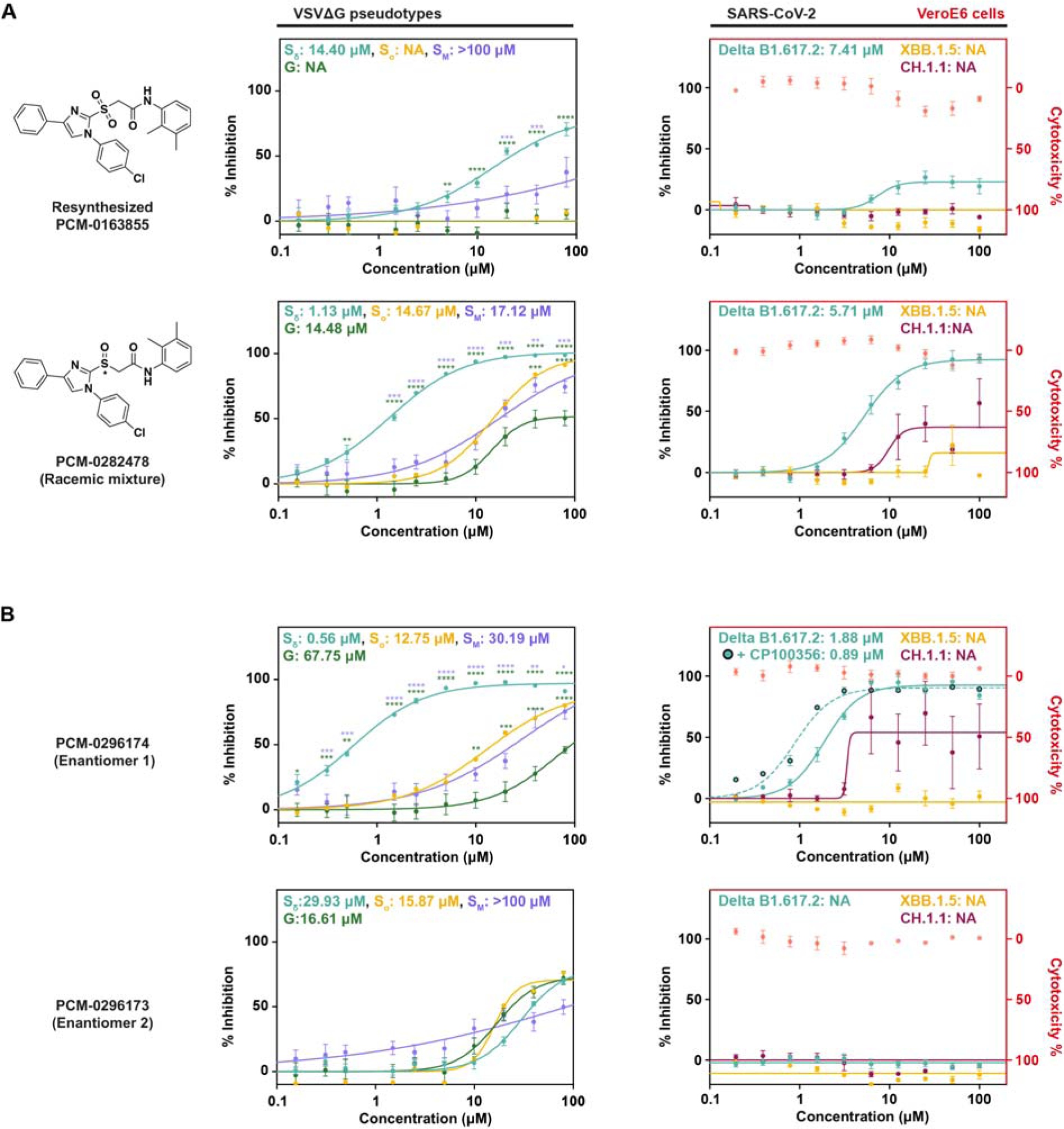
Validation of resynthesized PCM-0163855 and its sulfoxide derivatives. **(A; Left)** Structures of PCM-0163855 and its sulfoxide derivative PCM-0282478, and **(Middle)** corresponding dose-response plots comparing the inhibitory activity against VSVΔG-S_O_, VSVΔG-S_ο_, VSVΔG-S_M_ or VSVΔG-G, showing that PCM-0282478 exhibits a broader selectivity for inhibition compared to PCM-0163855, which was inactive against VSVΔG-G and, after resourcing, against VSVΔG-S_M_ at all concentrations in pseudovirus based assay. **(Right)** The corresponding cytotoxicity profiles and dose-response plots comparing the inhibitory activity of *bona fide* SARS-CoV-2 variants, Delta B1.617.2, XBB.1.5 or CH.1.1, viral replication in Vero E6 cells. **(B; Left)** Dose-response plots of two enantiomers of PCM-0282478 comparing the inhibitory activity against VSVΔG-S_O_, VSVΔG-S_ο_, VSVΔG-S_M_ or VSVΔG-G, showing that only one enantiomer exhibits a broader selectivity for inhibition in pseudovirus based assay. **(Right)** The corresponding cytotoxicity profiles and dose-response plots comparing the inhibitory activity of *bona fide* SARS-CoV-2 variants. The active enantiomer, PCM-0296174 was also tested against Delta.B1.617.2 in presence of the multidrug resistance protein 1 (MDR1) inhibitor, CP100356. (A and B) Error bars represent the SEM. The statistical significance of the inhibitions was also determined. P: *≤0.05, **≤0.01, ***≤0.001 ****≤0.0001 (multiple unpaired t-tests comparing group means, accounting for individual variance in each concentration and pseudovirus). N(experiments)≥2, n(readings) ≥6.

PCM-0282478 was synthesized as a racemic mixture. To evaluate the contribution of each enantiomer, we separated them to PCM-0296173 and PCM-0296174, and evaluated their IC_50_ value against VSVΔG-S_O_, VSVΔG-S_ο_, VSVΔG-G, and VSVΔG-S_M_ **(Fig. 5 B)**. Only PCM-0296174 potently inhibited VSVΔG-S_O_ and was selective against VSVΔG-S_ο_ and VSVΔG-S_M_ (12.75 μM and 30.19 μM respectively). The IC_50_ of value of PCM-0296174 against VSVΔG-S_O_ was approximately half compared to the racemic mixture PCM-0282478 (0.56 μM vs 1.13 μM respectively), showing potency increased two-fold. Both PCM-0296173 and PCM-0296174 did not inhibit VSVΔG-G up to 10 μM. Further, only PCM-0296174 inhibited SARS-CoV-2 Delta (B.1.617.2) replication, and its IC_50_ improved by 2-folds in presence of an inhibitor against the multidrug resistance protein 1 (MDR1) efflux transporter (1.88 μM vs 0.89 μM respectively; **Fig. 5 B**). Conversely, PCM-0296174 inconsistently inhibited the omicron variant CH.1.1 and did not inhibit XBB.1.5 (**Fig. 5 B**). These experiments taken together, suggest that PCM-0296174 is a cell permeable selective S glycoprotein inhibitor.

## Discussion

To identify potential CoVs fusion inhibitors, we developed an HTS platform that relies on a phenotypic assay of infection using well-characterized, replication-deficient VSVΔG pseudoviruses that can be studied at biosafety level 2 (BSL-2) and express a fluorescent reporter upon infection. This allowed us to produce and accurately titer pseudoviruses featuring the S glycoproteins from SARS-CoV-2 variants and MERS-CoV, which was essential for establishing pseudoviruses preparation with titers of 10^4^ infections per ml (**Fig. 1 F**). If the titers had dropped below 10^3^ infections per ml, the feasibility of using 384-well plates would have been compromised, rendering the screening process both economically and logistically prohibitive. We employed fluorescent reporters, instead of the more commonly used luciferase, demonstrating their advantages which include direct measurement of single infection events, high signal-to-noise ratio, and reduced processing. Moreover, fluorescent reporters with different wavelengths enable the simultaneous examination of multiple viruses under the same experimental conditions (**Fig. 1 G**).

We leveraged the HTS platform to screen a comprehensive library of approximately 200,000 compounds, targeting potential CoV fusion inhibitors (**Fig. 3 A**). Our three-tiered assay incorporated two class I viral glycoproteins from phylogenetically distant CoVs (SARS-CoV-2 & MERS-CoV), and a class III glycoprotein from VSV^57,58^ (**Fig. 3 A**). The primary screen against VSVΔG-S_W_ yielded 733 compounds with inhibitory activity, including known inhibitors of proteases, ubiquitin-specific peptidases, and viral gene expression regulated by the HSP90 protein family. The secondary screen showed that most of these compounds also inhibit VSVΔG-G, yielding only 65 putative Spike-specific inhibitors. Finally, the tertiary screen against VSVΔG-S_M_ separated receptor binding and putative S2-specific inhibitors, highlighting four compounds, and the known TMPRSS2 inhibitor Nafamostat (**Fig. 3 C-D**).

Following validation only PCM-0163855 was retained as a potential selective S glycoprotein inhibitor **(Fig. 4)** and its sulfoxide derivative PCM-0282478 inhibited VSVΔG-S_W_, VSVΔG-S_o_, VSVΔG-S_M_ and *bona fide* SARS-CoV-2 delta virus replication (**Fig. 5**). Finally, we showed that only one of the PCM-0282478 derived enantiomers, PCM-0296174, inhibits VSVΔG-S_W_, VSVΔG-S_o_, VSVΔG-S_M_ and *bona fide* SARS-CoV-2 delta virus replication, demonstrating its potential as a broad-spectrum inhibitor of CoVs. Note that we observed the Omicron CH.1.1 variant yielded an IC_50_ value with higher variability and no inhibitory activity against the XBB.1.5 variant (**Fig. 5**). This may suggest that the unique mutations in the XBB.1.5 variant may alter the binding site for PCM-0296174 on the Spike protein. Furthermore, given the likely poor solubility of PCM-0296174 (indicated by a calculated log D of 4), it is possible that we are underestimating the calculated IC_50_ values for variants that are well inhibited and cannot calculate it for the CH.1.1 variant. Synthesizing more soluble and active derivatives of PCM-0296174 will likely help clarify the breadth of selectivity and mode of action. Intriguingly, Nafamostat failed to inhibit *bona fide* SARS-CoV-2 delta virus replication, perhaps due to the capacity of CoVs to infect cells in a TMPRSS2-independent pathway (**Fig. S4**).

Our findings reinforce the utility of the HTS platform in identifying novel CoVs inhibitors with the potential to deepen our understanding of coronavirus biology. It also highlights the significance of rigorous compound triage, which is instrumental in averting the dissemination of ambiguous results. The discovery of PCM-0296174, a completely new compound that we synthesized and separated from the racemic mixture, as a promising compound with broad-spectrum antiviral will surely catalyze future research. Hence, the present study lays the groundwork for potential development of a new class of small molecules, holding promise for mitigating the impacts of future pandemics.

## Methods

### DNA constructs

pCAGGS-G, encoding the Vesicular Stomatitis Virus G glycoprotein from the Indiana serotype (VSV-G), was a kind gift from Benjamin Podbilewicz (Technion - Israel Institute of Technology)^59^. pcDNA3.1-SARS2-Spike-C9, encoding the Spike glycoprotein of Wuhan SARS-CoV-2 fused to a C-terminal C9 tag (S_W_) was a kind gift from Fang Li (Addgene plasmid # 145032; http://n2t.net/addgene:145032; RRID:Addgene_145032)^60^. pCG1-SARS2-Spike-HA, encoding the Wuhan SARS-CoV-2 Spike protein fused to a C-terminal HA tag was a kind gift from Gideon Schreiber, pCMV3-SARS2-Spike-Flag, encoding the Wuhan SARS-CoV-2 Spike protein fused to a C-terminal FLAG tag, pCMV3-SARS2-SpikeΔ19, encoding the Wuhan SARS-CoV-2 Spike protein with 19 amino acids removed at the cytoplasmic tail, and pCMV3-SARS2-SpikeΔ19-Flag, with an added C-terminal FLAG tag were a kind gift from and Yosef Shaul (Weizmann Institute of Science)^61^. pcDNA3.1-SARS-CoV-2-S_α_, SARS-CoV-2-S_δ_, and SARS-CoV-2-S_ο,_ encoding the Spike glycoprotein of Wuhan SARS-CoV-2 variants fused to a C-terminal C9 tag were generated by DNA synthesis (GeneScript). pcDNA3.1-MERS-Spike-C9, encoding the Spike glycoprotein of the MERS-CoV (S_M_) fused to a C-terminal C9 tag was generated by sub-cloning the MERS Spike protein from a pLVX-EF1alpha-MERS-Spike plasmid (Weizmann Plasmid Bank) into a pcDNA3.1 expression plasmid using the GeneArt Gibson Assembly HiFi master mix (Thermo Fisher Scientific cat. no. A46627). The following primers were used to generate the vector fragment (Primers V1: CGCACAAGGTCCACGTCCACGGCTCCACCGAGACATCCC and V2: AGAAAAACTGAATGAATCATGCTAGCCAGCTTGGGTC; template DNA: pcDNA3.1-SARS-Spike alpha) and the insert fragment (Primers I1: GGAGACCCAAGCTGGC TAGCATGATTCATTCAGTTTTTCTGCTCATGTTTC and I2: TGGGATGTCTCGGTGGAGCCGTGGACGTGGACCTTGTGC; template DNA: pLVX-EF1alpha-MERS-Spike).

### Cell culture

Baby Hamster Kidney cells (BHK-21; ATCC) were maintained in Dulbecco’s modified Eagle medium (DMEM, Gibco, USA) supplemented with 10% fetal bovine serum (FBS, Biological Industries, Israel), 1% penicillin-streptomycin (PS, Biological Industries) and 25mM HEPES (Biological Industries).

Human Embryonic Kidney-293T cells (HEK-293T; ATCC), HEK-293T overexpressing either ACE2 (HEK-293T-ACE2) or both ACE2 and TMPRSS2 (HEK-293T-ACE2-TMPRSS2), or ACE2, DPP4 and TMPRSS2 (HEK-293T-ACE2-TMPRSS2-DPP4) were cultured in DMEM supplemented with 8.1% FBS, 1% PS, 25mM HEPES (Biological Industries). HEK-293T-ACE2, HEK-293T-ACE2-TMPRSS2, and HEK-293T-ACE2-DPP4-TMPRSS2 were maintained by supplementing the culture medium with 1 μg/ml Puromycin (Sigma-Aldrich, USA), and 1.5 μg/ml Blasticidin (InvivoGen, USA) respectively. All cell lines were cultured in a 5% CO2 incubator at 37°C. HEK-293T-ACE2 and HEK-293T-ACE2-TMPRSS2 were a kind gifts from Yosef Shaul (Weizmann Institute of Science)^61^. HEK-293T-ACE2-TMPRSS2-DPP4 cells were established by lentiviral transduction. 0.3 x 10^6^ HEK-293T-ACE2-TMPRSS2 cells were seeded in each well of a 6 well plate and cultured to 70-80% confluency. The growth medium was replaced with 1 ml of medium containing human CD26/DPP4 pre-packaged lentiviral particles (LTV-CD26, G&P Biosciences) at a multiplicity of infection (MOI) of 5. To enhance transduction efficiency, 1:1000 of polybrene Infection Reagent (Sigma-Aldrich) was added to the medium. The cells were incubated with the lentivirus for 24h at 37 °C with 5% CO2. After the transduction period, the viral supernatant was removed, and a fresh growth medium containing 1 μg/ml Puromycin was added to the cells to initiate the selection process.

### Preparation of VSV**Δ**G pseudoviruses

To generate 5 ml of glycoprotein X complemented pseudoviruses (VSVΔG-X), 1.2×10^6^ BHK-21 cells were plated in a 100 mm dish one day prior to transfection. The cells were transfected at 75-80% confluence with 5 μg of a plasmid expressing the viral glycoprotein. 24 h after transfection, cells were infected with VSVΔG-G pseudovirus at a MOI of 5, with 1:1000 of polybrene Infection Reagent (Sigma-Aldrich). Cells were incubated for 1 h at 37 °C in a 5% CO_2_ incubator shaking every 15 min. Post infection, cells were washed with DPBS six times, and the medium was replaced with a 5 ml growth medium. 30 h post-infection, cells and the supernatant containing the pseudoviruses were collected and centrifuged at 500 g for 10 mins at 4°C. Clean supernatant was collected, aliquoted, and frozen at −80 °C for further experiments.

### Viral titer

To determine viral titers of VSVΔG complemented pseudoviruses, 10 μl of pseudovirus suspension was subsequently dispensed in a 384 well plate (Greiner, Austria) in sextuplicate. 10K/20 μl HEK-293T-ACE2-TMPRSS2 cells were added to each well and incubated for 15 mins at RT to allow the cells to settle. To maximize infection, assay plates were then subjected to 1000g centrifugation for 1 h at RT and incubated for 24 h in a 5% CO_2_ incubator at 37 °C.

After 24 hours, the plates were imaged using cell discoverer 7 (Carl Zeiss, Germany) in widefield mode with sCMOS 702 camera (Carl Zeiss, Germany). Images were acquired using a ZEISS Plan-APOCHROMAT 5x / 0.35 Autocorr Objective. ZEN blue software 3.1 (Carl Zeiss, Germany) was used for image acquisition using 470 nm excitation for the acquisition of the infected channel. The infected cells were segmented and counted using cellpose^62^. Infection/ml were extrapolated by calculating the geometric mean of the number of infected cells per well (each containing 10 μl) multiplied by 100. Where applicable the pseudovirus containing supernatant was first incubated with anti-G neutralizing antibody (1:1000 dilution, clone 1E9F9) for 1 h at room temperature (RT) to remove residual VSVΔG-G background infections.

### Compound libraries

The compound collection of the Nancy and Stephen Grand Israel National Center for Personalized Medicine (G-INCPM) was used for screening (https://g-incpm.weizmann.ac.il/units/WohlDrugDiscovery/chemical-libraries). 173,227 unique compounds from commercial sub-collections were used. The composition of the screening set was 0.7% Bioactive collections (Selleck Chemicals, USA), 7.6% HitFinder (Maybridge, USA), 10.8% Drug Like Set (Enamine, Ukraine), 26.8% DiversetCL (Chembridge, USA), 54.1% Diversity (ChemDiv, USA). Compounds were stored in 100% DMSO in acoustic dispenser certified plates. Hit compounds were purchased from Sigma-Aldrich, Aldrich Market Select (Sigma-Aldrich), Enamine (Ukraine) or MolPort (Latvia) chemical suppliers.

### Phenotypic assay: pseudotype-based HTS imaging inhibition assay

Compounds were spotted using Echo 555 Liquid Handler (Beckman Coulter, USA) on 384-well assay plates in 10 μM final concentration. To avoid background from any residual VSVΔG-G activity, pseudoviruses were incubated with anti-G neutralizing antibody (1:5000, clone 1E9F9) for 1 h at RT. Subsequently, 10 μl of the pseudovirus suspension were dispensed using Multidrop™ Combi (Thermo Fisher Scientific, USA) and incubated with compounds for 15 mins at RT. HEK-293T-ACE2-TMPRSS2 were trypsinized, counted and diluted to 0.5×10^6^ cells/ml. 20 μl of this cell suspension was dispensed (MultidropCombi) in each well containing the compound and the neutralized pseudoviruses and incubated for an additional 15 mins at RT. To maximize infection, assay plates were then subjected to 1000g centrifugation for 1 h at RT and incubated overnight in a 5% CO_2_ incubator at 37°C.

To account for the toxicity at 10 μM for a compound, wells were stained for nuclei with 5 μg/ml Hoechst 33342 (Thermo Fisher Scientific, USA) and incubated for 10 mins at 37 °C, 5% CO_2_ before live cell imaging was done by (ImageXpressMicro-confocal, Molecular Devices, USA) equipped with 4x S Fluor lens in two channels: filter set DAPI (ex 377 nm/em 447 nm) and FITC (ex 475 nm/em 536 nm) for total cells and infected cells, respectively.

Images were analyzed using MetaXpress CME (Molecular Devices, USA) to quantify the number of total and infected cells. Settings for segmentation: cell/nuclei size 5-30 μm and intensity >2000AU.

### Dose response assay

For dose-response assay the compounds were serially diluted to cover a range of 40-0.31 μM. The assay is identical to phenotypic assay as described above. Six different VSVΔG viruses representing G, WT, Alpha, Delta, Omicron, and MERS were tested. Data were deposited in CDD vault (Collaborative Drug Discovery platform), and dose response curves were analyzed from image analysis. Dose response curves were generated and fitted to the Levenberg– Marquardt algorithm that is used to fit a Hill equation to dose-response data.

### Cell viability assay

To assess compounds toxicity to cells, a copy of the compound as in dose-response experiment were assayed for live-dead assay. Cells were stained with Hoechst (as above) in addition to 1.5 μM Propidium Iodide (Life Technologies. Cat. P3566) and 2 μM Calcein AM (Life Technologies. Cat. C3099). Image analysis was performed by MetaXpress adjusted to quantify total, live or dead cells.

### Liquid Chromatography-Mass Spectrometry (LC/MS)

Flash chromatography was performed by automated CombiFlash® Systems (Teledyne ISCO, USA) with RediSep Rf Normal-phase silica gel columns (Teledyne ISCO) or Silica gel Kieselgel 60 (0.04-0.06 mm) columns (Merck, USA). Purification of the compounds was performed using preparative HPLC; Waters Prep 2545 Preparative Chromatography System, with UV/Vis detector 2489, using XBridge® Prep C18 10μm 10×250 mm Column (PN: 186003891, SN:161I3608512502). Reaction progress and compounds purity was monitored by Waters UPLC-MS system: Acquity UPLC® H class with PDA detector, ELSD detector, and using Acquity UPLC® BEH C18 1.7 μm 2.1×50 mm Column (PN:186002350, SN 02703533825836). MS-system: Waters, SQ detector 2. UPLC Method: 5 min gradient 95:5 Water: Acetonitrile 0.05% formic acid to Acetonitrile 0.05% formic acid, flowrate 0.5 mL/min, column temp 40°C.

### Synthesis of PCM-0163855 and PCM-0282478

All reagents, solvents and building blocks used for the synthesis were purchased from Sigma-Aldrich, Merck, Acros Organics (USA), Tzamal D-Chem Laboratories (Israel), Enamine, Combi-Blocks (USA) and MolPort chemical Suppliers and used for synthesis without further purification. All solvents used for reactions were of HPLC grade. Solvent and reagent abbreviations: Ethyl acetate (EtOAc), Dichlormethane (DCM), Dimethylformamide (DMF), 1,8-Diazabicyclo[5.4.0]undec-7-ene (DBU), Diisopropylethyl amine (DIPEA), Trifluoroacetic acid (TFA). Reactions on microwave were done on Biotage Initiator+ (Biotage, Sweden). ^1^H NMR spectra were recorded on a Bruker Avance III −300 MHz, 400 MHz and 500 MHz spectrometer, equipped with QNP probe. Chemical shifts are reported in ppm on the δ scale and are calibrated according to the deuterated solvents. All *J* values are given in Hertz.

*Ethyl 2-((1-(4-chlorophenyl)-4-phenyl-1H-imidazol-2-yl)thio)acetate **(2)***: To a 5 mL crimp vial, ethyl bromo acetate (87.3 mg, 0.52 mmol) was added followed by 1-(4-chlorophenyl)-4-phenyl-1H-imidazole-2-thiol ***(1)***^63^ (100.0 mg, 0.35 mmol) and DIPEA (182.0 µl, 1.05 mmol). The vial was crimped and heated at 90 °C for 5min in microwave reactor, and the reaction was cooled and diluted with EtOAc (10 ml), the organic layer was washed 1X water, 1X brine and dried on Na_2_SO_4_. The organic layer was concentrated onto silica (0.5 g) and purified on a Combi-Flash Systems (Teledyne ISCO), using a 24 g silica gel column gradient (15 min) from DCM to EtOAc. The fraction that eluted in 50% EtOAc gave compound **2** (123.0 mg, 94 % yield).

*2-((1-(4-chlorophenyl)-4-phenyl-1H-imidazol-2-yl)thio)acetic acid **(3)***: Compound **2** (123.0 mg, 0.33 mmol) was dissolved in THF (2 ml), then LiOH (197.0 mg, 8.24 mmol) was dissolved in water (2 ml), and the freshly prepared solution was added to the reaction and stirred overnight. The reaction mixture was cooled to 0 °C and acidified to pH = 4 with HCl 1M (∼ 9 ml), the aqueous layer was extracted 3x EtOAc, the combined organic layers were washed with brine dried on Na_2_SO_4_. The organic layer was concentrated to give compound **3** (106.2 mg, 93% yield).

*2-((1-(4-chlorophenyl)-4-phenyl-1H-imidazol-2-yl)thio)-N-(2,3-dimethylphenyl)acetamide **(4)***: In a 25 mL round bottom flask, compound **3** (106.0 mg, 0.31 mmol) and 2,3-dimethylaniline (72.7 mg, 0.46 mmol) was dissolved in DMF (2 ml). Then DIPEA (214.0 µl, 1.23 mmol) was added followed by HATU (128.6 mg, 0.34 mmol) and the reaction was stirred for 12h. The reaction was then poured into brine (10 ml) and the aqueous layer was extracted 3 X EtOAc, the combined organic layer was washed with 1x water, 1x brine and dried on Na2SO4. The organic layer was concentrated onto 0.5 g silica and purified on Combi-Flash Systems, using a 24 g silica gel column gradient (17 min) from DCM to EtOAc, the fraction that eluted in 50% EtOAc gave compound **4** (108.2 mg, 79% yield).

*2-((1-(4-chlorophenyl)-4-phenyl-1H-imidazol-2-yl)sulfonyl)-N-(2,3-dimethylphenyl)acetamide **(PCM-0163855)***: To a 25 mL round bottom flask, compound **4** (19.5 mg, 0.044 mmol) was dissolved in DCM (2 ml), then mCPBA (24.4 mg, 0.11 mmol) was added in one portion. After 2h an additional amount of mCPBA (10 mg) was added and the reaction was stirred overnight. The reaction mixture was quenched with sodium sulfite (30 µl) and concentrated on rotary evaporator. Purification of the crude reaction mixture by HPLC water to MeCN gradient 45 min, the desired compound eluted in 80% MeCN to give **PCM-0163855** (13.5 mg, 62% yield). 1H-NMR (300 MHz, DMSO-d6) δ 9.82 (s, 1H), 8.22 (s, 1H), 7.88 (d, 2H, *J* = 8 Hz), 7.60-7.50 (m, 4H), 7.48-7.40 (m, 2H), 7.37-7.29 (m, 1H), 7.08-7.00 (m, 3H), 4.67 (s, 2H), 2.24 (s, 3H), 1.97 (s, 3H); ES-LRMS: (*m/z*) calculated for C_25_H_23_ClN_3_O_3_S ([M+H]+) 480.1, found 480.3.

*2-((1-(4-chlorophenyl)-4-phenyl-1H-imidazol-2-yl)sulfinyl)-N-(2,3-dimethylphenyl)acetamide **(PCM-0282478)***: To a 25 ml round bottom flask, compound **4** (52.0 mg, 0.012 mmol) was dissolved in DCM (2 ml), then mCPBA (28.6 mg, 0.013 mmol) was added in one portion and the reaction was stirred overnight. The reaction mixture was quenched with sodium sulfite (30 µl) and the reaction was concentrated on rotary evaporator. Purification of the crude reaction mixture by HPLC water to MeCN gradient 45 min, the desired compound eluted in 80% MeCN to give **PCM-0282478** (35.1 mg, 65% yield). Chiral Separation was performed at Lotus Separations (USA, http://lotussep.com/). 1H-NMR (300 MHz, DMSO-d6) δ 10.02 (s, 1H), 8.31 (s, 1H), 7.95 (d, 2H, *J*=7.5 Hz), 7.78-7.68 (m, 4H), 7.51-7.41 (m, 2H), 7.38-7.30 (m, 1H), 7.06-6.98 (m, 3H), 4.98 (d, 1H, *J*=14 Hz), 4.64 (d, 1H, *J*=14 Hz), 2.21 (s, 3H), 1.96 (s, 3H); (ES-LRMS: (*m/z*) calculated for C_25_H_23_ClN_3_O_2_S ([M+H]+) 464.1, found 464.3.

### SARS-CoV-2 replication assay

Clear-bottomed 384-well black plates were seeded with 3,000 Vero E6 cells or Vero TMPRSS2 cells per well. The following day, individual compounds were added at ten specified concentrations, 2 h prior to infection. Each plate included DMSO (0.5 %) and remdesivir (25 µM; SelleckChem) controls. After the pre-incubation period, the cells were exposed to the Delta B.1.617.2 inoculum (at a multiplicity of infection of 0.05 PFU/Vero E6 cell and 0.2 PFU/Vero TMPRSS2 cell). After a one-hour adsorption at 37°C, the supernatant was aspirated and replaced with 2% FBS/DMEM media containing the respective compounds at the indicated concentrations. The cells were then incubated at 37 °C for 2 days. Supernatants were collected and heat inactivated at 80 °C for 20 minutes. The Luna Universal One-Step RT-qPCR Kit (New England Biolabs) was used for the detection of viral genomes in the heat-inactivated samples performed through reverse transcription quantitative polymerase chain reaction (RT-qPCR). Specific primers targeting the N gene region of SARS-CoV-2 (5′-TAATCAGACAAGGAACTGATTA-3′ and 5′-CGAAGGTGTGACTTCCATG-3′) were utilized. The cycling conditions involved an initial step at 55 °C for 10 minutes, followed by 95 °C for 1 minute. Subsequently, 40 cycles were carried out with denaturation at 95 °C for 10 seconds and annealing/extension at 60 °C for 1 minute using an Applied Biosystems QuantStudio 6 thermocycler. The quantity of viral genomes is expressed as Ct and was normalized against the Ct values of the negative and positive controls.

In parallel, cytotoxicity was assessed using the CellTiter-Glo luminescent cell viability kit (Promega). 3,000 cells/well of Vero E6 were seeded in white with clear bottom 384-well plates. The following day, compounds were added at concentrations indicated. DMSO only (0,5%) and 10 μM camptotecin (Sigma-Aldrich) controls were added in each plate. After 48 h incubation, 10 μl of CellTiter Glo reagent was added in each well and the luminescence was recorded using a luminometer (Berthold Technologies) with 0.5 sec integration time.

Raw data were normalized against appropriate negative (0 %) and positive controls (100 %) and are expressed in % of viral replication inhibition or % of cytotoxicity. Curve fits and IC_50_/CC_50_ values were obtained in Prism using the variable Hill slope model.

Delta B.1.617.2: The variant was supplied by Virus and Immunity Unit in Institut Pasteur headed by Olivier Schwartz. It was isolated from a nasopharyngeal swab of a hospitalized patient who had returned from India. The swab was provided and sequenced by the Laboratoire de Virologie of the Hopital Européen Georges Pompidou (Assistance Publique des Hôpitaux de Paris). XBB.1.5: The strain hCoV-19/France/PDL-IPP58867/2022 was supplied by the National Reference Centre for Respiratory Viruses hosted by Institut Pasteur (Paris, France) and headed by Dr. Etienne Simon-Lorière. The human sample from which strain hCoV-19/France/PDL-IPP58867/2022 was isolated has been provided from the Centre Hospitalier de Laval. CH.1.1: The strain hCoV-19/France/NAQ-IPP58166/2022 was supplied by the National Reference Centre for Respiratory Viruses hosted by Institut Pasteur (Paris, France) and headed by Dr. Etienne Simon-Lorière. The human sample from which strain hCoV-19/France/NAQ-IPP58166/2022 was isolated has been provided from the Selas Cerballiance Charentes.

### Statistical analysis and tools

Data from analyzed images was processed using Genedata Screener (Genedata, Switzerland). Normalization was the percent of infection where neutral control is 0% inhibition of infection and “No-Virus” control is 100% inhibition of infection. Further chemoinformatic data visualizations were made with Certara D360 software, which is integrated with the CDD database. Excel (Microsoft, USA) was used to analyze the data and Prism (GraphPad, USA) was used to plot the data. Whenever comparing between two conditions, data was analyzed with two tailed student’s t-test. Measurements are reported as mean of at least three biological repeats, and the error bars denote standard error of mean (SEM). Throughout the study, threshold for statistical significance was considered for p-values≤0.05, denoted by one asterisk (∗), two (∗∗) if P≤0.01, three (∗∗∗) if P <0.001 and four (∗∗∗∗) if P≤0.001.

## Acknowledgements

We thank members of the Avinoam laboratory, and The Wohl Drug Discovery Institute of the Nancy and Stephen Grand Israel National Center for Personalized Medicine (G-INCPM) for discussions. We also thank Yosef Shaul, Romano Strobelt and Julia Adler for cell lines and plasmids, and Benjamin Podbilewicz, Clari Valansi and Gideon Schreiber for plasmids, Galit Cohen and Shirly Valter for assisting with compound plating. This research was supported by Israel Science Foundation (grant no. 3729/20), Bina Nurture Program, and the European Research Council (ERC) Proof of Concept Grant (101100758 – Inhibicov). OA also acknowledges funding from the Henry Chanoch Krenter Institute for Biomedical Imaging and Genomics, the Schwartz Reisman Collaborative Science Program, and the European Research Council (ERC) under the European Union’s Horizon 2020 research and innovation program (grant agreement no 851080). OA is an incumbent of the Miriam Berman presidential development chair. This research was also supported in part by funding from the INSTITUT PASTEUR, Research Applications and Industrial Relations Department (DARRI) and the ANRS-MIE (BIOVAR and PRI projects of the EMERGEN research program) to FA.

## Author contributions

OA, SK, and EOP conceived the study, designed, and executed the experiments with contributions from YEA and NS. SK, NK, HMB, and OA performed and analyzed the screen. KS performed the mass spectroscopy experiments, compound purification and synthesis. JC and FA supervised all experiments performed under BSL-3 conditions, and JC, EG, J.T-R performed experiments with SARS-CoV-2 viruses. SK and OA wrote the manuscript with inputs from all authors.

---

**Fig. S1.**
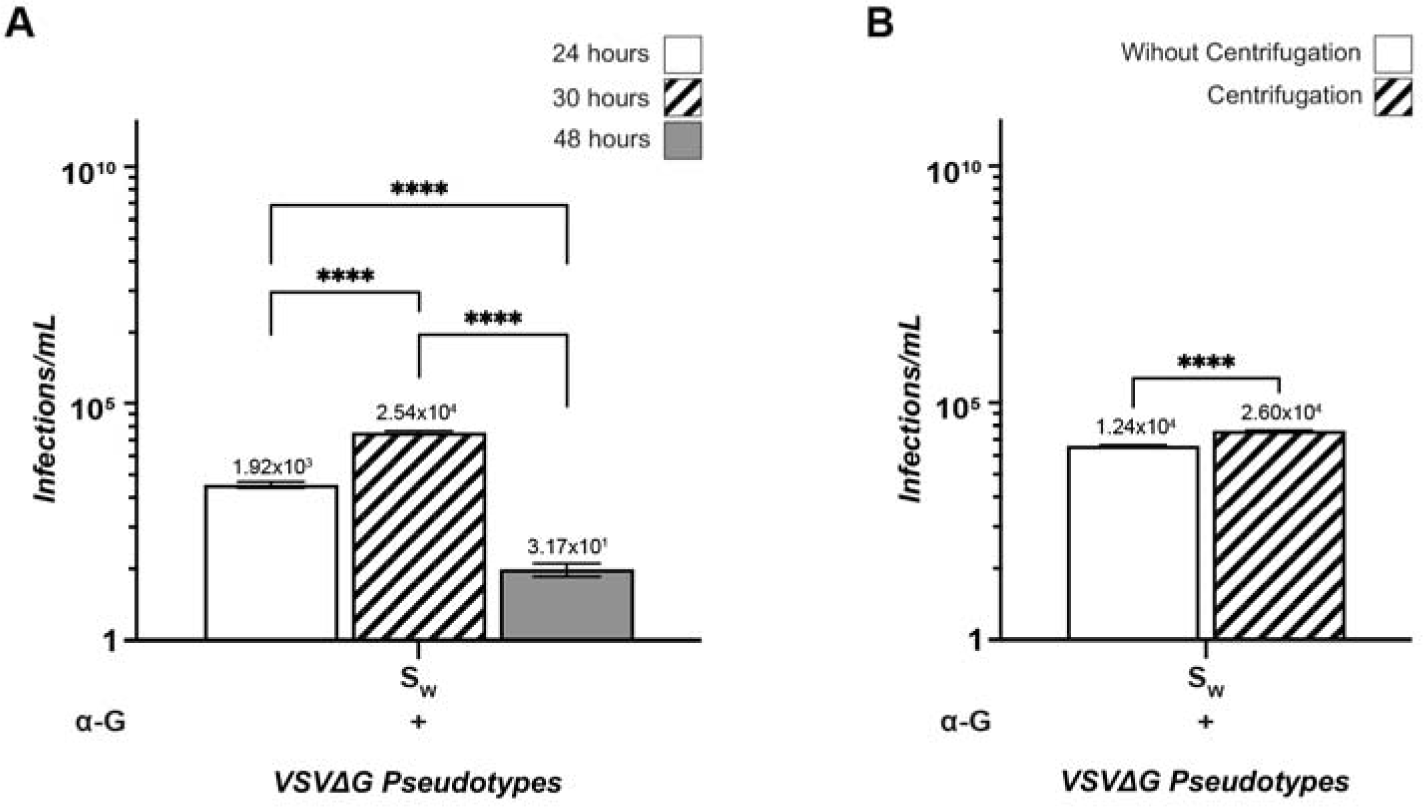
Optimization of pseudovirus titer. **(A)** Infections/ml of VSVΔG-S_W_ pseudoviruses present in the harvested supernatant at different times, indicating optimal titers at 30 hours post-harvest. **(B)** Infections/ml of samples subjected to centrifugation, indicating a two-fold increase in viral titer with centrifugation. Experiments were performed in the presence of a VSV-G neutralizing antibody to exclude any residual infection from VSVΔG-G that was left over from the production. The statistical significance of conditions was also determined. P: ****≤0.0001 (two-tailed unpaired t-tests). N(experiments)=3, n(readings)=9.

**Fig. S2.**
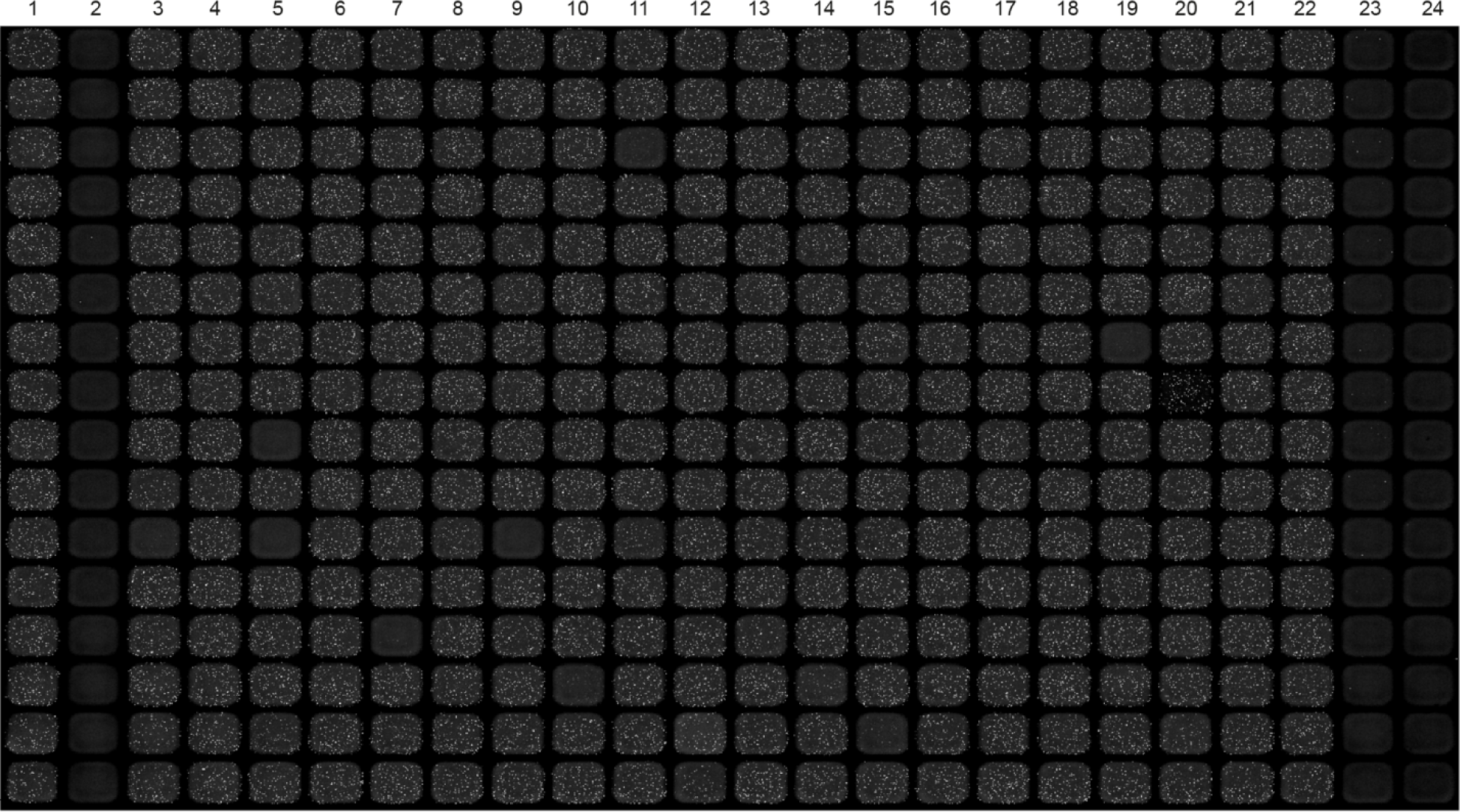
Overview of a 384-well plate from the screen. Column 1, the neutral controls, indicates 100% infection, and Column 2, the positive controls, signifies 0% infection or 100% inhibition. Columns 3-22 are spotted with compounds. Columns 23 and 24 are plated with VSVΔG-G pseudoviruses. An antibody against VSV-G (α-G) is added in column 23. All columns are supplemented with DMSO to achieve a final concentration of 0.01%.

**Fig. S3.**
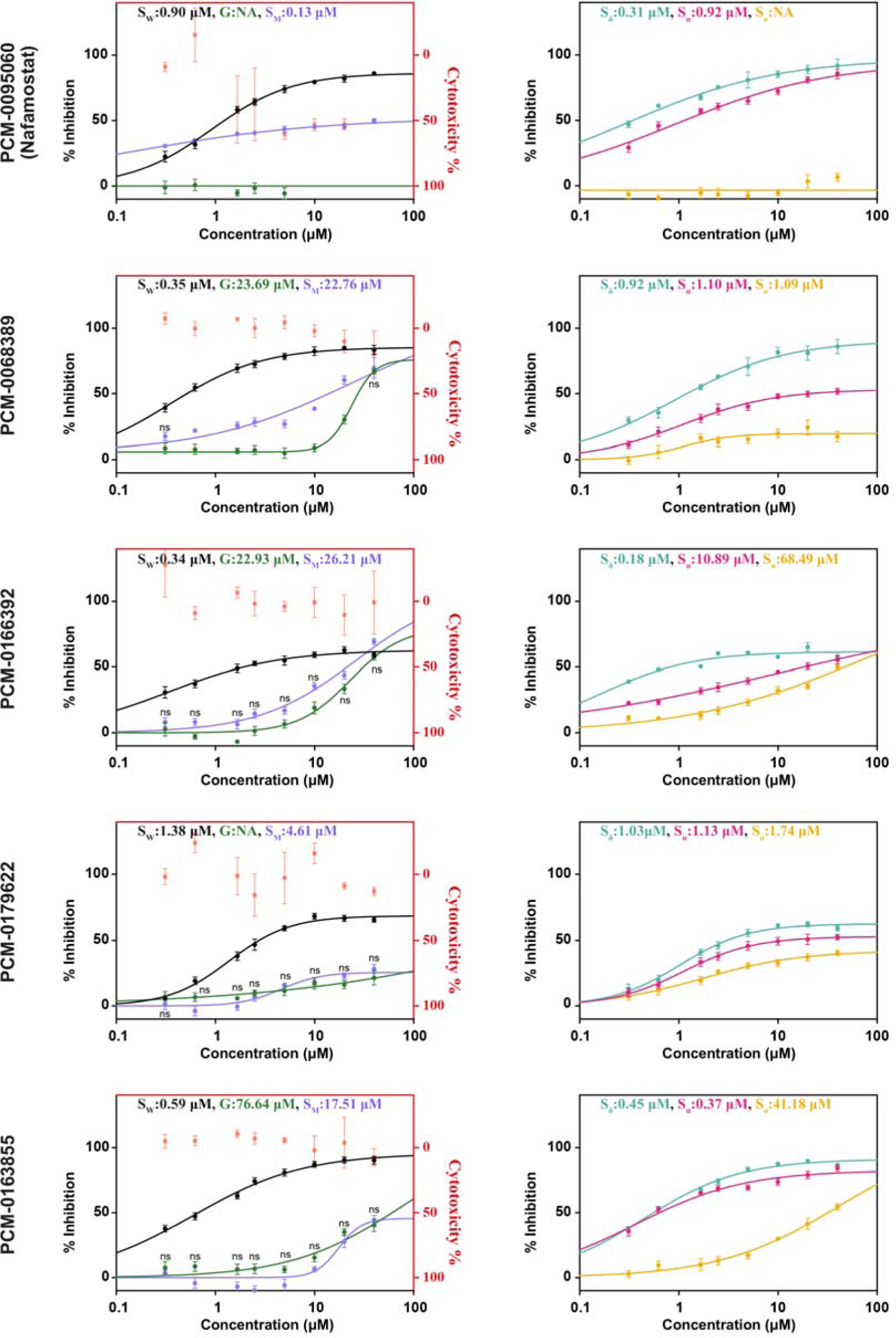
Dose-response activity and cytotoxicity of compounds before HPLC. (Left) Dose-response plot of the hits against VSVΔG-S_W_, VSVΔG-S_M_ or VSVΔG-G and their cytotoxity profile. (Right) Dose-response plots of hits against pseudoviruses with glycoproteins of SARS-CoV-2 variants (VSVΔG-S_α_, VSVΔG-S_O_, VSVΔG-S_ο_). Error bars represent the SEM. ns are the readings where there is no statistically significant difference between VSVΔG-S_M_ and VSVΔG-G at a given concentration. For all other readings, the P<0.05 (two-tailed unpaired t-tests).

**Fig. S4.**
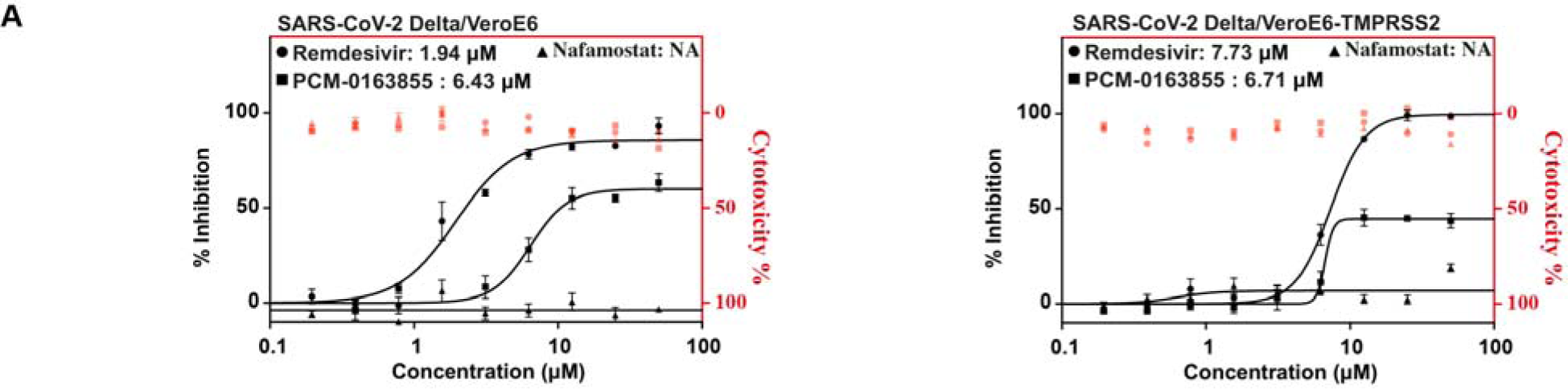
Validation of PCM-0163855 against *bona fide* SARS-CoV-2. **(A)** Cytotoxicity profile of PCM-0163855 and dose-response plots comparing the inhibitory activity of PCM-0163855, and known inhibitors Nafamostat and Remdesivir on SARS-CoV-2 delta variant on viral replication in Vero E6 cells with (left) and without (right) TMPRSS2 over expression.

**Fig. S5.**
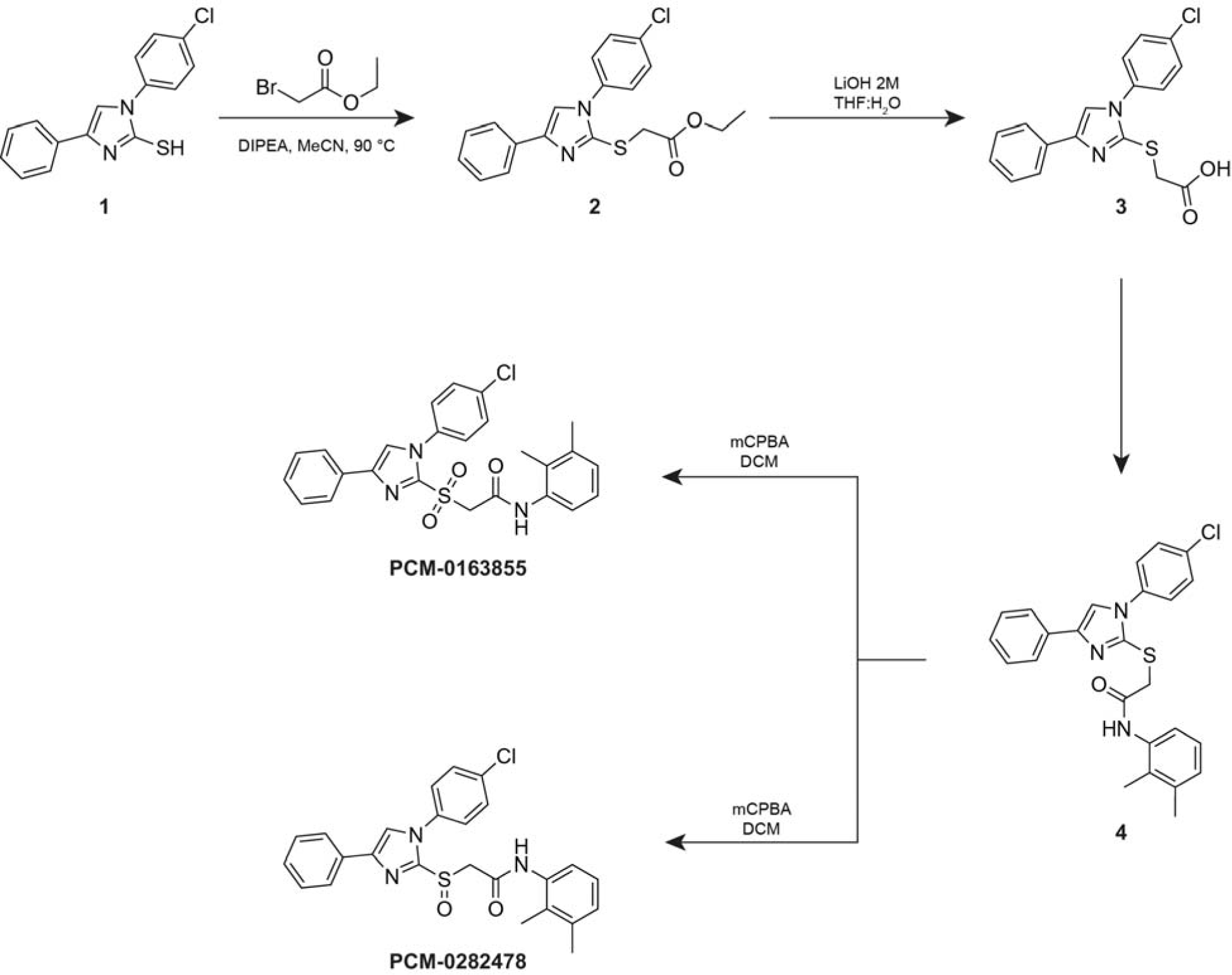
General reaction scheme for synthesis of PCM-0163855 and PCM-0282478.

**Fig. S6.**
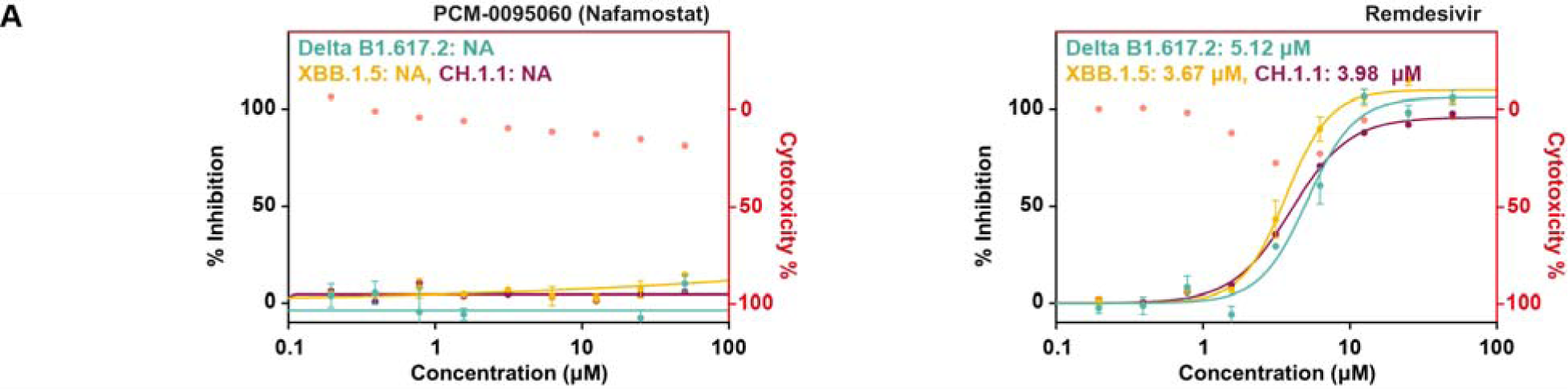
Activity of controls against *bona fide* SARS-CoV-2 variants. **(A)** Cytotoxicity profile of Nafamostat (left) and Remdesivir (right) against SARS-CoV-2 variants, Delta B1.617.2, XBB.1.5 and CH.1.1, on viral replication in Vero E6 cells.

**Table 1.**
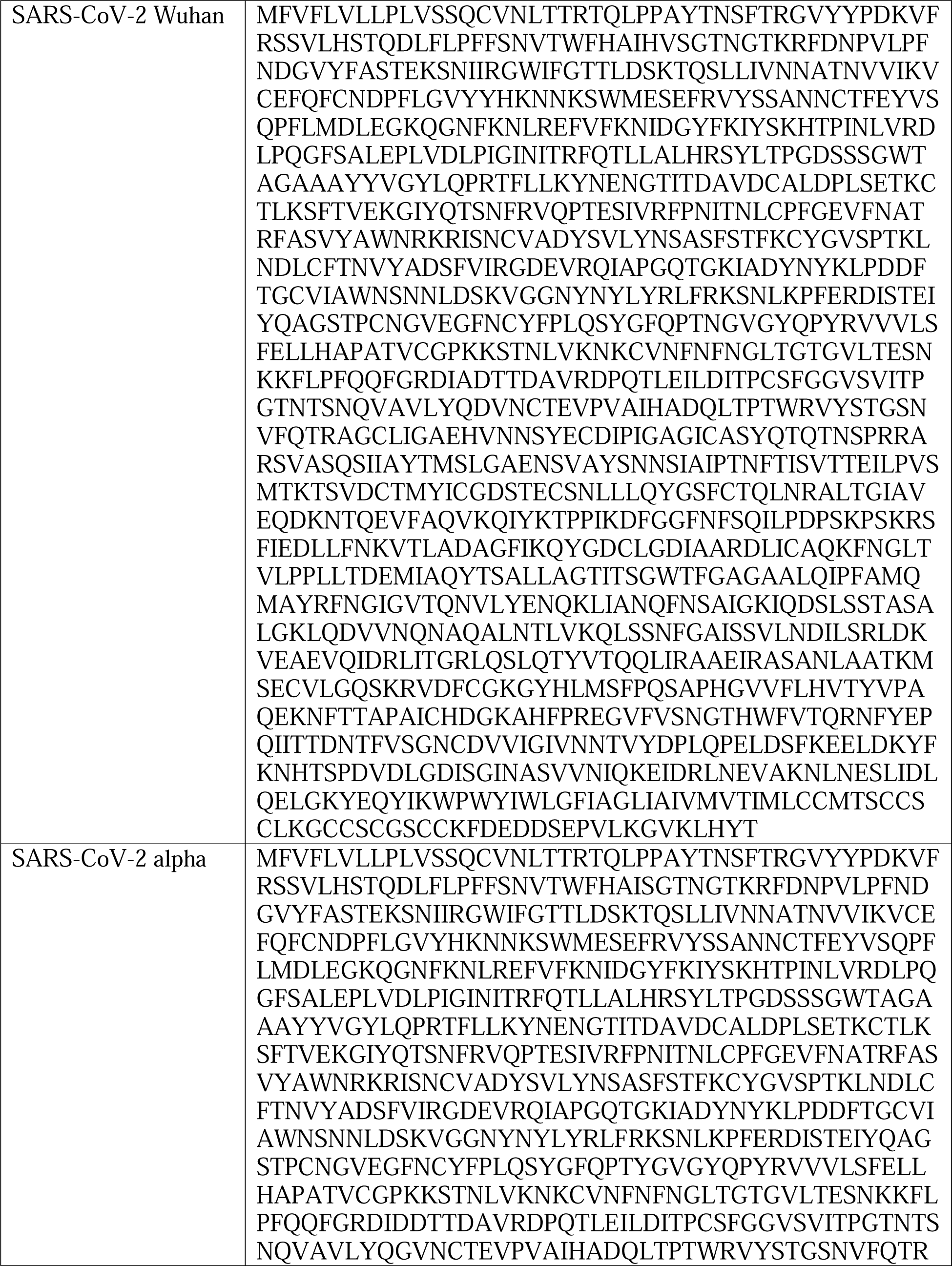
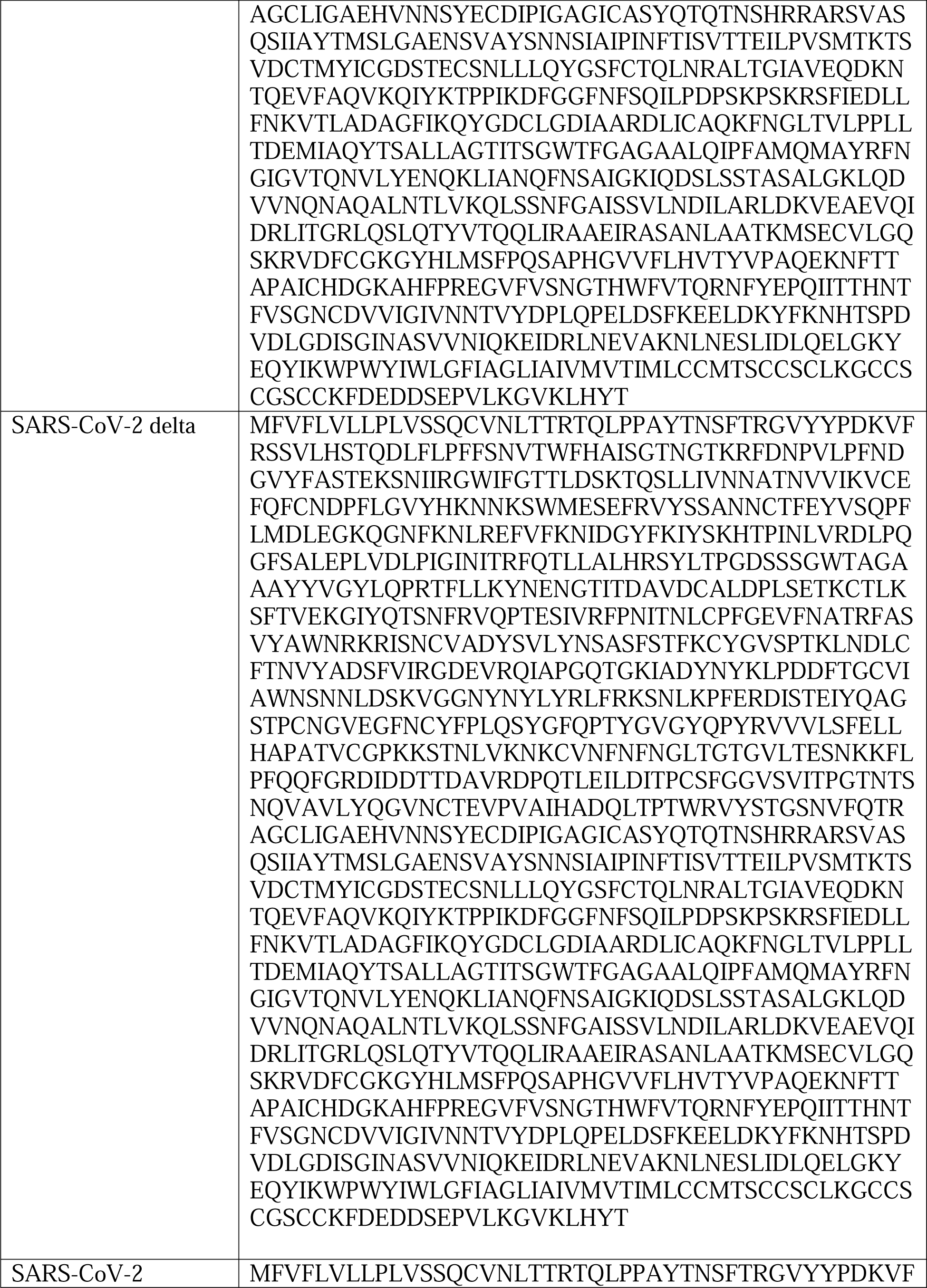
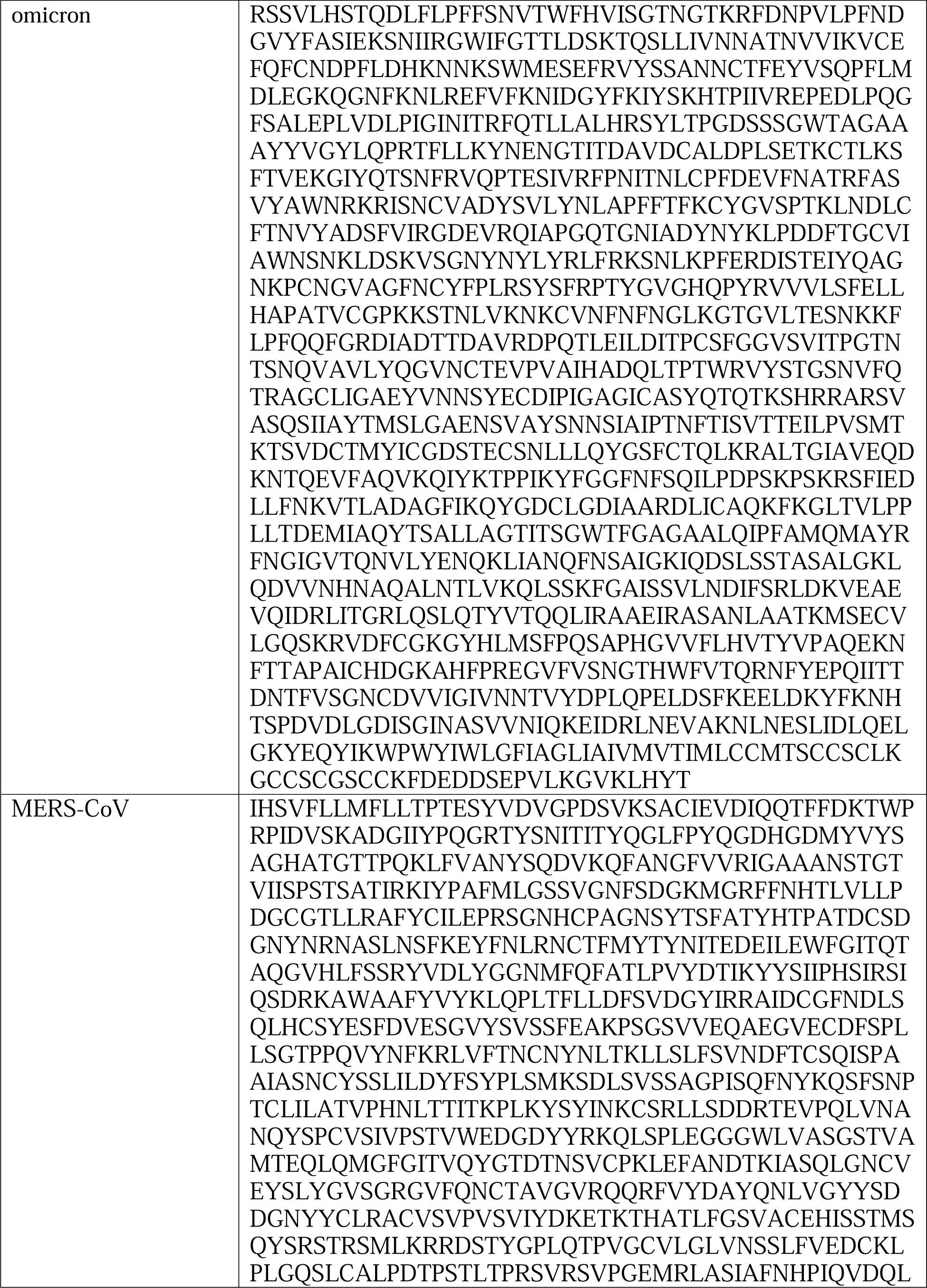
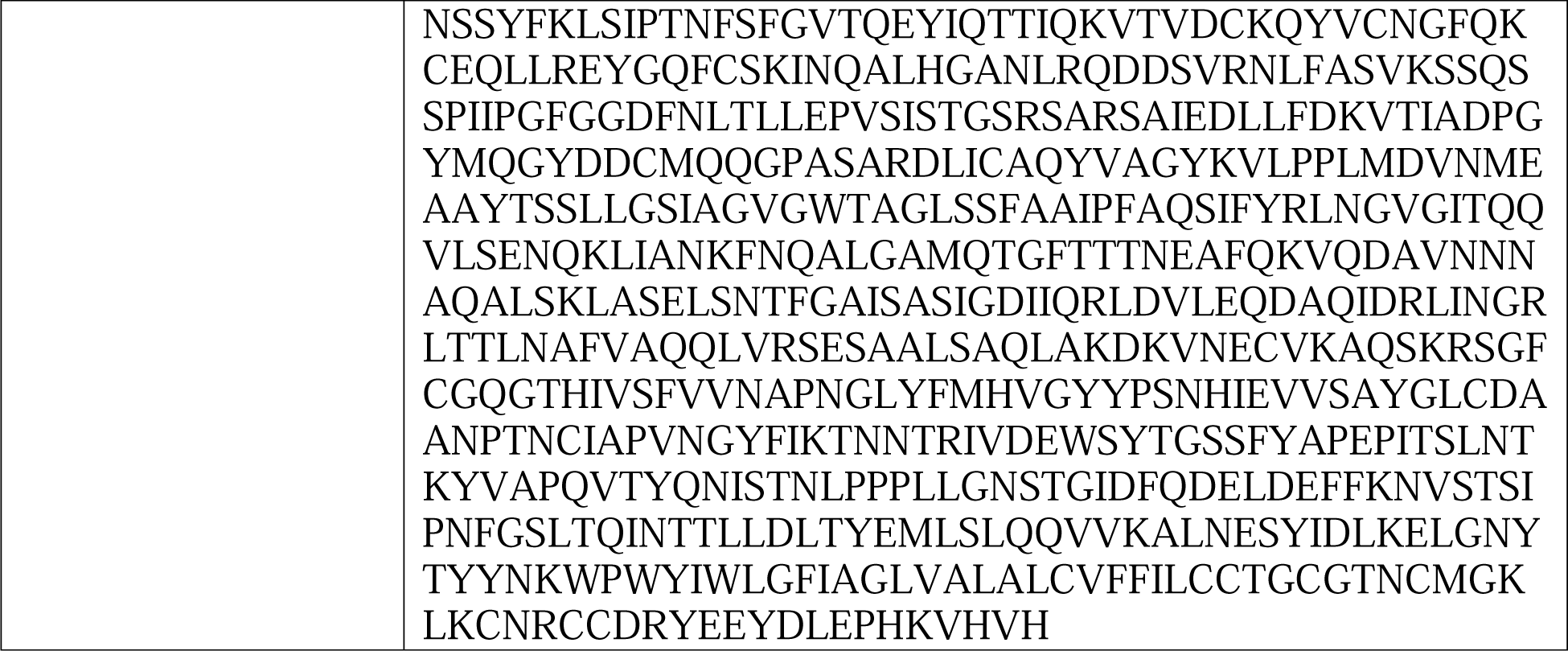
Sequences of Coronavirus Spike proteins used in this study.

